# Notch signaling and taxis mechanims regulate early stage angiogenesis: A mathematical and computational model

**DOI:** 10.1101/569897

**Authors:** Rocío Vega, Manuel Carretero, Rui D.M. Travasso, Luis L. Bonilla

## Abstract

During angiogenesis, new blood vessels sprout and grow from existing ones. This process plays a crucial role in organ development and repair, in wound healing and in numerous pathological processes such as cancer progression or diabetes. We present here a mathematical model of early stage angiogenesis that permits to explore the relative importance of mechanical, chemical and cellular cues. Endothelial cells proliferate and move over an extracellular matrix by following external gradients of Vessel Endothelial Growth Factor, adhesion and stiffness, which are incorporated to a Cellular Potts model with a finite element description of elasticity. The dynamics of Notch signaling involving Delta-4 and Jagged-1 ligands determines tip cell selection and vessel branching. Through their production rates, competing Jagged-Notch and Delta-Notch dynamics determine the influence of lateral inhibition and lateral induction on the selection of cellular phenotypes, branching of blood vessels, anastomosis (fusion of blood vessels) and angiogenesis velocity. Anastomosis may be favored or impeded depending on the mechanical configuration of strain vectors in the ECM near tip cells. Numerical simulations demonstrate that increasing Jagged production results in pathological vasculatures with thinner and more abundant vessels, which can be compensated by augmenting the production of Delta ligands.

**Author Summary:** Angiogenesis is the process by which new blood vessels grow from existing ones. This process plays a crucial role in organ development, in wound healing and in numerous pathological processes such as cancer growth or in diabetes. Angiogenesis is a complex, multi-step and well regulated process where biochemistry and physics are intertwined: with signaling in vessel cells being driven by both chemical and mechanical mechanisms that result in vascular cell movement, deformation and proliferation. Mathematical models have the ability to bring together these mechanisms in order to explore their relative relevance in vessel growth. In this work, we present a mathematical model of early stage angiogenesis that is able to explore the role of biochemical signaling and tissue mechanics. We use this model to unravel the regulating role of Jagged, Notch and Delta dynamics in vascular cells. These membrane proteins have an important part in determining the leading cell in each neo-vascular sprout. Numerical simulations demonstrate that increasing Jagged production results in pathological vasculatures with thinner and more abundant vessels, which can be compensated by augmenting the production of Delta ligands.

## Introduction

Angiogenesis is a process by which new blood vessels sprout and grow from existing ones. This ubiquitous phenomenon in health and disease of higher organisms [1], plays a crucial role in the natural processes of organ growth and repair [2], wound healing [3], or inflammation [4]. Angiogenesis imbalance contributes to numerous malignant, inflammatory, ischaemic, infectious, and immune diseases [2,5], such as cancer [6–9], rheumatoid arthritis [10], neovascular age-related macular degeneration [11], endometriosis [12, 13], and diabetes [14].

Either when a tissue is in hypoxia or during (chronic or non-chronic) inflammation, cells are able to activate signaling pathways that lead to the secretion of pro-angiogenic proteins. The Vascular Endothelial Growth Factor (VEGF) is one of these proteins and it is necessary and sufficient to trigger angiogenesis. Present in different isoforms, VEGF diffuses in the tissue, and is able to bind to extracellular matrix (ECM) components (its binding affinity is different for distinct VEGF isoforms), forming a well defined spatial concentration gradient in the direction of increasing hypoxia [15, 16]. When the VEGF molecules reach an existing vessel, they promote the dwindling of the adhesion between vessel cells and the growth of newer vessel sprouts. VEGF also activates the tip cell phenotype in the vessel endothelial cells (ECs) [17]. The tip cells grow filopodia rich in VEGF receptors, pull the other ECs, open a pathway in the ECM, lead the new sprouts, and migrate in the direction of increasing VEGF concentration [18]. Branching of new sprouts occur as a result of crosstalk between neighboring ECs [19].

As the new sprouts grow, ECs have to alter their shape to form a lumen connected to the initial vessel that is capable of carrying blood [20–24]. Moreover, in order for the blood to be able to circulate inside the new vessels, the growing sprouts have to merge either with each other or with existing functional mature vessels [25]. The process by which sprouts meet and merge is called anastomosis [25–29].

Nascent sprouts are then covered by pericytes and smooth muscle cells, which provide strength and allow vessel perfusion. Poorly perfused vessels may become thinner and their ECs, in a process that inverts angiogenesis, may retract to neighboring vessels leading to more robust blood circulation [30, 31]. Thus, the vascular plexus remodels into a highly organized and hierarchical vascular network of larger vessels ramifying into smaller ones [32].

Angiogenesis is therefore a multi-step, complex and well regulated process where biochemistry and physics are intertwined; with signaling in ECs being driven by both chemical and mechanical mechanisms that result in EC proliferation, mechanical deformation and cell movement.

In particular, the dynamical and biochemical processes that take place at the tip of every growing sprout are determinant for the growth, morphology and function of the resulting neo-vasculature. When an EC has the tip cell phenotype (which is triggered by the binding of VEGF to VEGF Receptor 2, VEGFR2) its membrane becomes rich in Delta-4 transmembrane proteins [19, 33]. These proteins bind the Notch transmembrane proteins in the neighboring cells triggering the Notch signaling pathway. The activation of this pathway down-regulates VEGFR2 and Delta-4, forcing the neighboring cells not to be in the tip cell phenotype, and to acquire the stalk cell phenotype [34]. Stalk ECs are characterised by a higher proliferation rate [18] triggered by both VEGF and by the tension exerted on them by the tip cell [35]. The sprouts are able to grow due to proliferation of the stalk ECs.

The ECs can interchange dynamically their phenotypes from tip to stalk. In fact, in the growing sprout the stalk ECs behind the tip cell are often able to overtake the tip cell and to take its place, thereby becoming tip cells and driving sprout elongation [36, 37]. This dynamic behavior ensures that there is always a cell at the front of the sprout with the tip phenotype capable of exerting a contractile force on the matrix, degrading and remodelling matrix fibres and opening a pathway for the sprout to grow.

EC metabolism is strongly connected with this cycling dynamics at the tip of sprouts [38], and it is determinant to vascular patterning, pruning and sprouting [39–41]. The ability of ECs to rearrange themselves is essential for vessel remodelling [30]. Moreover, this dynamics at the tip is only possible due to the regulation of VE-cadherin expression in ECs by the Notch signaling pathway [36, 42, 43]. When the Notch-driven tip-stalk pattern is absent (due to very high VEGF levels, for example) the EC rearrangement dynamics stops [43]. In that case the vessels become thicker and sprouting is severely hampered. Hence, the Notch signaling pathway is pivotal in determining the morphology of blood vessel networks.

Importantly, the dynamics of the ECs’ phenotypes in a growing sprout can be rather complex. While at moderate values of VEGF lateral inhibition by tip ECs can be observed [44], at higher VEGF concentrations the situation is different. Recently it has been experimentally observed that high levels of VEGF lead to synchronisation of phenotypes between cells at the sprout [45]. This phenomenon had first been suggested by theoretical models [17]. The model suggested that ECs in a sprout under high VEGF levels initiate acquiring the tip cell phenotype simultaneously, and then all simultaneously trigger the lateral inhibition by the Delta-Notch signaling, losing the tip phenotype and moving towards the stalk phenotype, only for the process to start again. Synchronised oscillatory behaviour in Delta-4 levels in EC cells has been observed under these conditions [45]. In this way, high VEGF hinders the symmetry breaking needed for the lateral inhibition to take place in the sprout.

Recent mathematical models of Notch signaling in angiogenesis have also predicted states where the cells in the sprout are in a third intermediate state and neither in the tip nor in the stalk phenotype [46, 47]. The Jagged-1 transmembrane protein is an important partner in the regulation of the Notch signaling in angiogenesis, and its introduction in the computational models permit to predict these intermediate EC phenotypes [47–50].

Jagged-1 is a ligand of Notch and competes with Delta-4 in angiogenesis [51]. Experiments have shown that when the lateral inhibition pattern induced by Delta-Notch signaling is present, the levels of Jagged-1 follow the EC phenotype: they are lower in tip cells and higher in stalk cells (contrary to what happens with the levels of Delta-4) [52]. However, ECs are able to control independently the levels of Jagged-1 (for example by reaction with proteins of the Fringe family [51]), and therefore they are able to control the sensitivity to Notch-mediated lateral inhibition. Moreover, Jagged-1 also plays an important role in making the Notch mechanism capable of lateral induction, whereby a stalk EC may induce its neighbors to acquire a phenotype equal to its own [53]. For these motives, it is extremely important to understand the implications of Jagged-1 levels in sprouting angiogenesis. Mathematical models should integrate the knowledge of Delta-Notch-Jagged signaling with the dynamics of EC organization in a sprout to better understand how the communication between ECs in angiogenesis is mediated by Jagged-1.

Numerous mathematical models of angiogenesis study the growth of blood vessels and irrigation using continuum methods, cellular automata, and hybrid methods [35, 54–73]. Cellular Potts Models (CPM) [74, 75] of angiogenesis and vasculogenesis have been particularly successful in capturing vascular cell shape [76], vascular structure [65, 77] and in integrating the role of extra-cellular matrix (ECM) mechanics and structure [66, 78–80] in the development of the vasculature.

Many of these models use simplified models of the Notch pathway to determine the separation between sprouts [25, 67, 69]. However, very detailed models of the Notch signaling pathway that integrate the dynamics of filopodia growth and of anastomosis have been developed [17, 81]. These detailed models also shed light into the regulation of VE-Cadherin by Delta-Notch [36, 43] and into the coupling between EC metabolism, Delta-Notch and EC rearrangement dynamics at the tip [38].

Moreover, cell based mathematical models that include Jagged-1 and Fringe have been developed in the contexts of cell differentiation [48–50] and angiogenesis [47]. However, these models of sprouting angiogenesis use a fixed geometry of a linear array of cells, without taking into account that ECs in a sprout are elongated and have many neighbors, and that they move and proliferate. Therefore, to describe the regulating effect of Jagged-1 in the sprouting dynamics we need to integrate dynamical models that take into account Jagged-1 with a CPM that takes into account cell shape and movement. In the present paper, we carry out this integration process for angiogenesis in the early stage, before sprouts form a lumen, become perfused and can regress. We use a CPM that incorporates cell motion following increasing gradients of VEGF (chemotaxis), of adhesion to substrate (haptotaxis) and of substrate stiffness (durotaxis), as well as cell proliferation and the Notch signaling pathway. This model will permit to explore the relative importance of mechanical, chemical and cellular cues in angiogenesis.

The section Mathematical Model describes the CPM coupled with the Delta-Notch-Jagged dynamics. In the section Results and Discussion, we present the results of the simulation and how Jagged-1 determines sprouting dynamics. Finally, in the last section we draw the conclusions of the manuscript.

## Mathematical model

The mathematical model consists of a CPM in which the dynamics of the Notch signaling pathway in endothelial cells selects tip and stalk ECs. Tip ECs move by chemotaxis, haptotaxis and durotaxis and stalk cells proliferate. Vessel branching and anastomosis appear as a result of combined cell signaling, mechanical and chemical taxis.

### Cellular Potts model

#### Square grid

We consider a square domain Ω of side *L* with grid points (*x_i_*, *y_j_*), where *x_i_* = *ih*, *y_j_* = *j h* with *i, j* = 0, … , *M* − 1, *h* = *L/*(*M* − 1), and *M* is the number of nodes on a side of the square. The square contains *M × M* grid points and (*M* − 1)^2^ elementary squares (pixels), each having an area *L*^2^*/*(*M* − 1)^2^. To enumerate nodes, we use left-to-right, bottom-to-top order, starting from node 0 on the bottom left corner of the square and ending at node *M*^2^ − 1 on the rightmost upper corner. In numerical simulations, we use *L* = 0.495 mm.

#### Objects, spins and Metropolis algorithm

Pixels **x** can belong to different objects Σ_*σ*_, namely ECs, and ECM. The field (called spin in a Potts model) *σ*(**x**) denotes the label of the object occupying pixel **x** [74]. Each given spin configuration for all the pixels in the domain has an associated energy *H*({*σ*(**x**)}) to be specified below. At each Monte Carlo time step (MCTS) *t*, we select randomly a pixel **x**, belonging to object Σ_*σ*_, and propose to copy its spin *σ*(**x**) to a neighboring (target) pixel **x**′ that does not belong to Σ_*σ*(**x**)_. The proposed change in the spin configuration (spin flip) changes the configuration energy by an amount Δ*H*|_*σ*(**x**)→*σ*(**x′**)_, and it is accepted with probability (Metropolis algorithm) [66, 74]

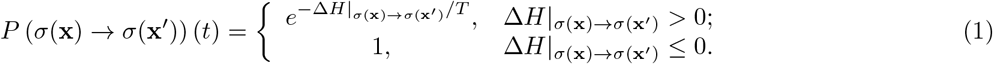

The temperature *T* > 0 is measured in units of energy and it is related to an overall system motility. We have selected *T* = 4 in our simulations. To obtain the equivalence between the number of MCTS and the time measured in experiments, we measured the pixel size in Fig. 1H of Ref [82], which is 0.9 *μ*m. According to Fig. 3C of the same reference, the vessel mean elongation is 150 pixels (135 *μ*m) in 36 hours for 50 ng/ml VEGF concentration. In our simulations, the vessel mean elongation is 495 *μ*m in 3001 MCTS. Thus, we set 1 MCTS to be 0.044 hours. This time for vessel growth also agrees with experiments in Ref. [83]. It is also possible to infer cellular forces from CPM simulations [84].

**Figure 1.**
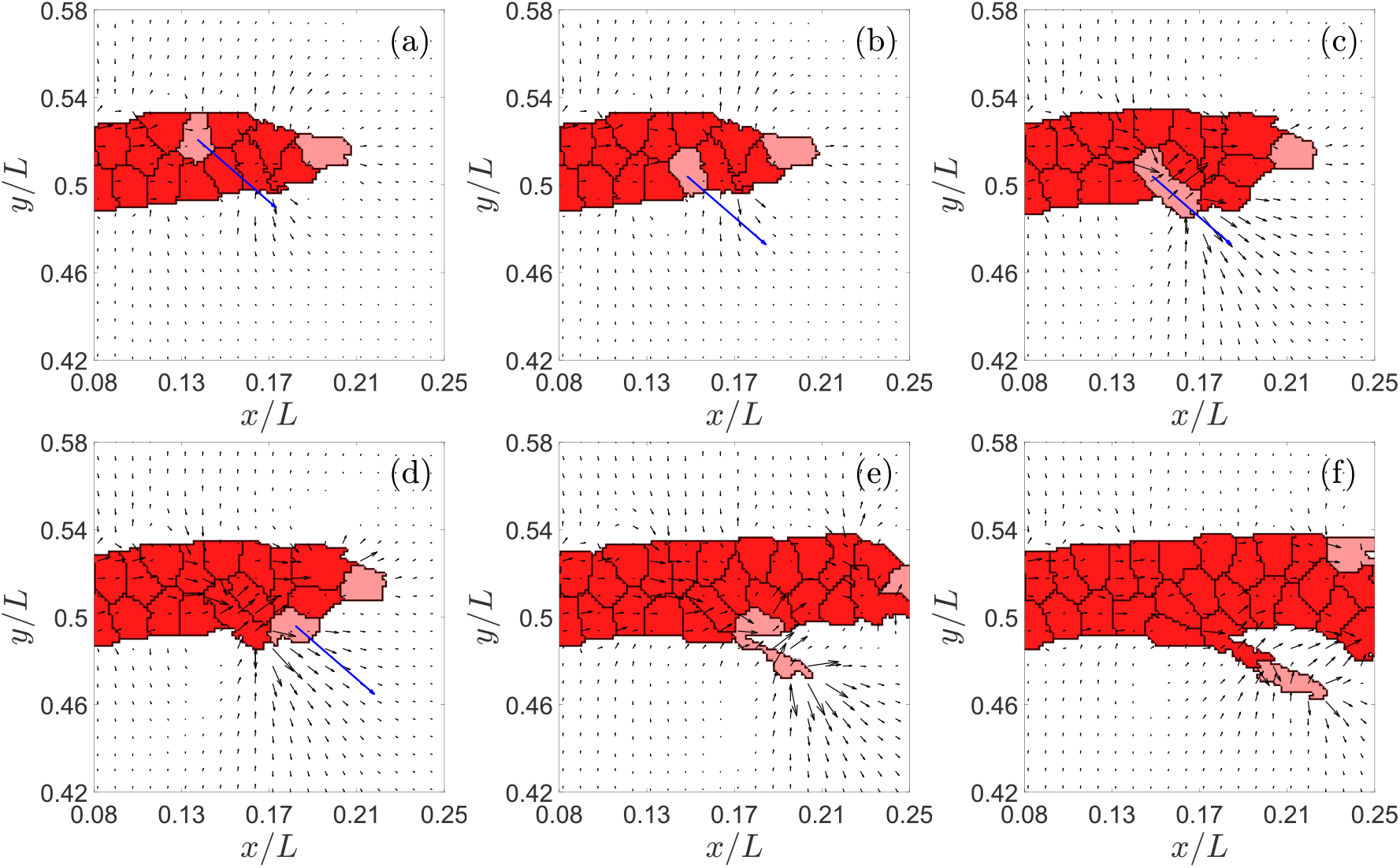
Example of tip cell exchange and branching in the direction of the blue arrow. Times in MCTS are: (a) 422, (b) 423, (c) 460, (d) 461, (e) 545, (f) 630. The black arrows in this figure represent the directions of largest eigenstrain and, therefore, they point to the likeliest direction of EC motion. The blue arrows indicate the actual direction of motion of a selected tip cell (marked in pink color) for the simulation we have carried out.

#### Energy functional

The energy functional *H* is

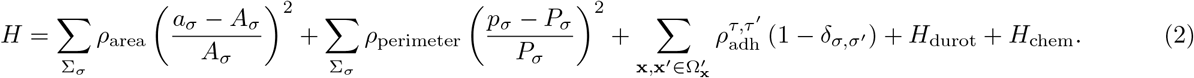

Here the two first terms are sums over cells and the third one sums over all pixels. We have

- *a_σ_* is the area of the cell Σ_*σ*_, *A_σ_* is the target area and *ρ*_area_ is the Potts parameter which regulates the fluctuations allowed around the target area. There are two cell types: non-proliferating tip and stalk cells with *A_σ_* = 78.50 *μ*m^2^ and proliferating cells with double target area, *A_σ_* = 157 *μ*m^2^. The target radius of a proliferating cell is 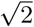 times that of a non-proliferating cell.
- *p_σ_* is the perimeter of the cell Σ*_σ_*, *P_σ_* is the target perimeter and *ρ*_perimeter_ is the Potts parameter which regulates the fluctuations allowed around the target perimeter. The target perimeters are *P_σ_* = 31.4 *μ*m for non-proliferating cells, and thrice this, *P_σ_* = 94.2 *μ*m, for proliferating cells.
- The Potts parameter 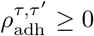 is the contact cost between two neighboring pixels. The value of this cost depends on the type of the object to which the pixels belong (cell or medium). Since *δ_σ,σ′_* is the Kronecker delta, pixels belonging to the same cell do not contribute a term to the adhesion energy.
- The net variation of the durotaxis term *H*_durot_ is [66]

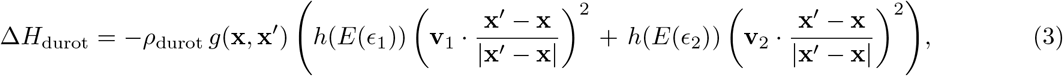

where *ρ*_durot_ is a Potts parameter, *g*(**x**, **x**′) = 1 for extensions and *g*(**x**, **x**′) = −1 for retractions, *ϵ*_1_ and *ϵ*_2_ and **v**_1_ and **v**_2_ (|**v**_1_| = |**v**_2_| = 1) are the eigenvalues and eigenvectors of the strain tensor *ϵ_T_*, respectively. They represent the principal strains and the strain orientation. *ϵ_T_* is the strain in the target pixel for extensions, and the strain in the source pixel for retractions. *h*(*E*) = 1/ (1 + exp (−*ω* (*E* − *E_θ_*))) is a sigmoid function with threshold stiffness *E_θ_* and steepness *ω*. *E*(*ϵ*) = *E*_0_ (1 + (*ϵ/ϵ*_st_)1_*ϵ*≥0_) is a function of the principal strains, in which *E*_0_ sets a base stiffness for the substrate, *ϵ*_st_ is a stiffening parameter and 1_*ϵ*≥0_ = {1, *ϵ* ≥ 0; 0, *ϵ* < 0}: strain stiffening of the substrate only occurs for substrate extension (*ϵ* ≥ 0), whereas compression (*ϵ* < 0) does not stiffen or soften the substrate. We have used the parameter values: *E_θ_* = 15 kPa, *E*_0_ = 10 kPa, *ω* = 0.5 kPa^−1^, and *ϵ*_st_ = 0.1 [66].
- The variation of the chemotaxis term *H*_chem_ is

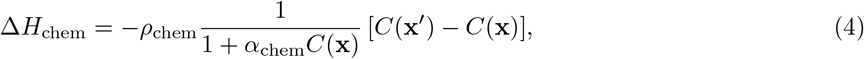

where *ρ*_chem_ ≥ 0 is a Potts parameter, *α*_chem_ = 0.3 and *C* is the VEGF concentration in the corresponding pixel.

The values of the Potts parameters are listed in Table 1. They are chosen according to those proposed by Bauer *et al.* [76] and Van Oers *et al.* [66] and adjusted so as to make that every term of the net variation of the hamiltonian have the same order. The perimeter contribution, absent in Refs. [66, 76], is small compared to the other terms in Eq. (2), so that it only affects the computations in extreme cases (e.g., extremely thin cells, thin cells that stick to the blood vessel). We have added a factor in the chemotaxis term to regulate the fluctuation around the resting VEGF concentration. Note that if *α*_chem_ is equal to zero, we recuperate the original term of Bauer *et al.* [76]. The proposed value, *α*_chem_ = 0.3, is small.

**Table 1.** Dimensionless Potts parameters.

| Parameter | $\rho_{\text{area}}$ | $\rho_{\text{perimeter}}$ | $\rho_{\text{durot}}$ | $\rho_{\text{chem}}$ | $\rho_{\text{adh}}^{\tau, \tau'} \text{ (cell-cell)}$ | $\rho_{\text{adh}}^{\tau, \tau'} \text{ (cell-ECM)}$ |
| --- | --- | --- | --- | --- | --- | --- |
| Value | 9000 | 250 | 25 | 60000 | 8.25 | 16.50 |

What is the effect of changing the numerical values of the Potts parameters? As said before, with the values in Table 1, every term of the net variation of the hamiltonian has the same order. Variations of 10% or smaller in Potts parameters do not change the outcome of the simulations. Variations larger than 10% with respect to those in Table 1 produce unrealistic effects, which are as follows.

- *ρ*_area_. Larger increments force cells to reach their target area faster, thereby increasing cell proliferation. The corresponding term becomes more important than the chemotaxis mechanism, which produces slower evolution of vessels toward the hypoxic zone and large clumps of cells in the vessels. Large reductions of this Potts parameter produce irregular cell proliferation and a much larger variety of cell sizes.
- *ρ*_perimenter_. Large increments produce round cells, whereas large reductions (up to *ρ*_perimenter_ = 0) creates extremely long and narrow cells stuck to the vessel sprout due to the now dominant effect of the adhesion term.
- *ρ*_durot_. This parameter produces qualitative changes only if it is ten times larger than in Table 1. In such a case, durotaxis overwhelms chemotaxis and the perimeter penalty, leading to cells following the stiffness gradients and sticking to each other, which create very irregular vessels.
- *ρ*_chem_. Larger increments make chemotaxis dominant. Then cells become bigger and elongated and sprouts extend more rapidly. Sometimes tip cells separate from their sprouts as chemotaxis dominates adhesion effects. Larger reductions produce rounder cells that do not polarize along a specific direction, and produce wider and slower sprouts.
- 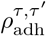. The adhesion Potts parameter take on different values for cell-cell and cell-ECM boundaries. If these values become equal (e.g., to 16.5), narrower sprouts are produced and there are cells that escape from them. Larger increments of cell-ECM adhesion, makes very costly for ECM to surround cells, which then stick to each other too much. Larger reductions of cell-ECM produces more elongated cells. Reducing cell-cell adhesion favors cells sticking to each other and acquiring irregular shapes since the zero energy for a pixel to be surrounded by other pixels of the same cell would be very similar to the small positive energy for the pixel to be surrounded by pixels of a different cell.

### Continuum fields at the extracellular scale

#### VEGF concentration

The VEGF concentration obeys the following initial-boundary value problem [76]:

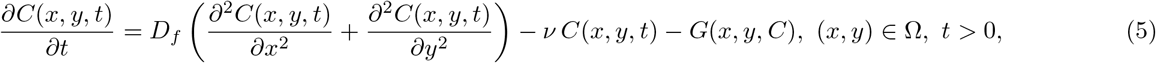

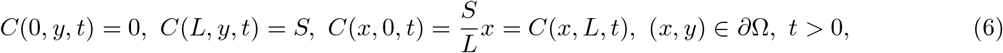

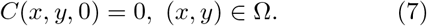

In Eq. (5), the amount of VEGF bound by an EC per unit time is

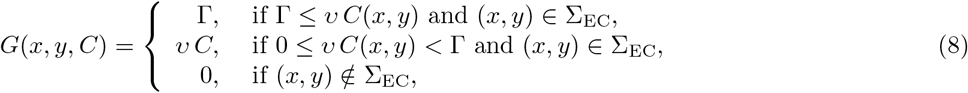

where *ν* = 1 h^−1^ and Γ = 0.02 pg/(*μ*m^2^ h) is the maximum amount of VEGF that it could be consumed by a cell per hour [62, 76]. Other values we use are *D_f_* = 0.036 mm^2^/h, *ν* = 0.6498/h, *S* = 0.0153 pg/*μ*m^2^ [76].

#### Strains

Following Ref. [66], we calculate the ECM strains by using the finite element method to solve the stationary Navier equations of linear elasticity:

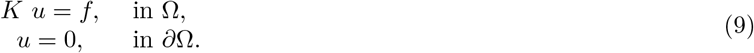

Here *K* is the stiffness matrix, *u* is the array of the *x* and *y* displacements of all nodes and *f* is the array of the traction forces per unit substrate thickness exerted by the cells. For nodes outside ECs, *f* = 0. For nodes inside ECs, each component 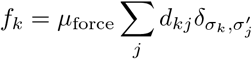 represents the traction stress on the *k*th node, *μ*_force_, times the sum of the distances, *d_kj_*, between the *k*th node and any node *j* in the same cell (*σ_k_* is the label of the cell at which node *k* belongs).

The global stiffness matrix *K* is assembled from the stiffness matrices *K_e_* of each pixel,

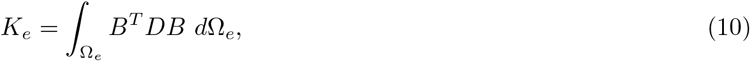

in which

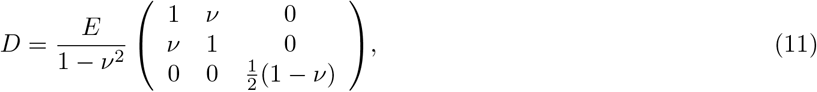

and *B* is the strain-displacement matrix for a four-noded quadrilateral pixel (finite element) [66]. *B* is a 3 × 8 matrix that relates the 8-component node displacement *u_e_* of each pixel to local strains *ϵ*,

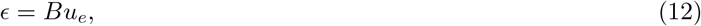

where *ϵ* = (*ϵ*_11_, *ϵ*_22_, *ϵ*_12_) is the 3-component column notation of the strain tensor

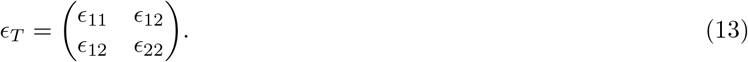

We have used the numerical values *E* = 10 kPa, *ν* = 0.45, and *μ*_force_ = 1 N/m^2^. With these definitions and the durotaxis term given by Eq. (3), ECs generate mechanical strains in the substrate, perceive a stiffening of the substate along the strain orientation, and extend preferentially on stiffer substrate. The simulated ECs spread out on stiff matrices, contract on soft matrices, and become elongated on matrices of intermediate stiffness [66].

### Signaling processes

The Notch signaling pathway is activated when Notch (transmembrane receptor) belonging to a particular cell interacts with Delta-4 or Jagged-1 (transmembrane ligands) belonging to its neighboring cell (trans-activation), thereby releasing the Notch intracellular domain (NICD). NICD then enters the nucleus and modulates the expression of many target genes of the Notch pathway, including both the ligands Delta and Jagged. However, when Notch of a cell interacts with Delta or Jagged belonging to the same cell, no NICD is produced; rather, both the receptor (Notch) and ligand (Delta or Jagged) are degraded (cis-inhibition) and therefore the signaling is not activated. For a given cell *i* surrounded by other cells, the equations describing this pathway are [47]

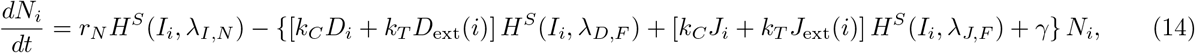

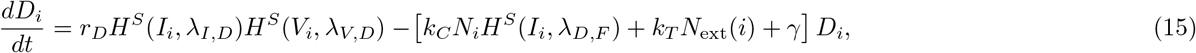

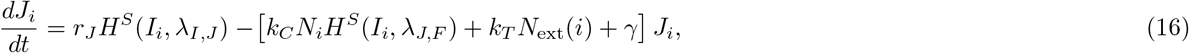

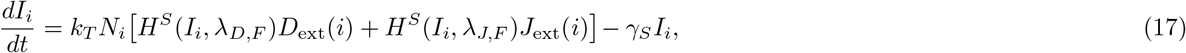

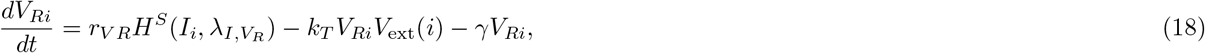

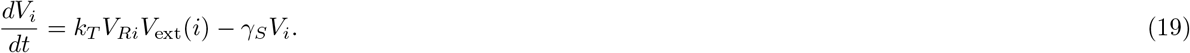

Here, *N_i_*, *D_i_*, and *J_i_* are the number of Notch, Delta-4, and Jagged-1 proteins in the *i*th cell, respectively, at time *t*. *I_i_*, *V_Ri_* and *V_i_* are the number of NICD, VEGF receptor and VEGF molecules, respectively, that are in the *i*th cell at time *t*. *r_N_*, *r_D_*, *r_J_*, and *r_V R_*, are the production rates of *N*, *D*, *J*, and *V_R_*, respectively. The cis-inhibition and trans-activation rates are *k_C_* and *k_T_*, respectively, whereas *γ* and *γ_S_* are degradation rates for *N*, *D*, *J*, *V_R_* and for *I*, *V*, respectively. These parameters, their representative values and units are listed in Table 2. All unknowns in Eqs. (14)-(19) are initially zero but changing these initial conditions does not alter the outcome of simulations.

**Table 2.** Rates appearing in Eqs. (14)-(19).

| Parameter | $r_N$ | $r_D, r_J, r_{V_R}$ | $k_C$ | $k_T$ | $\gamma$ | $\gamma_S$ |
| --- | --- | --- | --- | --- | --- | --- |
| Value | 1200 | 1000 | $5 \times 10^{-4}$ | $2.5 \times 10^{-5}$ | 0.1 | 0.5 |
| Unit | molec/h | molec/h | (h molec)$^{-1}$ | (h molec)$^{-1}$ | h$^{-1}$ | h$^{-1}$ |

Outside the *i*th cell, the number of *X* molecules is

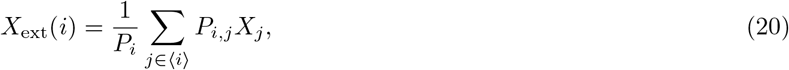

where *X* = *N, D, J*, and *j* ∈ ⟨*i*⟩ are the cells *j* sharing boundary of length *P_i,j_* with cell *i*. The perimeter of cell *i*, *P_i_*, minus ∑_*j*∈⟨*i*⟩_ *P*_*i,j*_ is the length of its boundary that is not shared with any other cell. Note that *X*_ext_(*i*) is simply the sum of all *X_j_* if the lengths *P_i,j_* are all equal and *P_i_* = ∑_*j*∈⟨*i*⟩_ *P*_*i,j*_ because the whole boundary of cell *i* is shared with other cells. As the cell moves and its boundaries fluctuate due to cellular Potts dynamics, the membrane protein levels of the neighboring cells interacting with the moving cell also vary. In this way, the production rates of the different proteins in a cell are directly influenced by the interactions with its neighborhood and, in particular, by the membrane fluctuations of the cell. *V*_ext_(*i*) is the number of VEGF molecules outside the *i*th cell that interact with VEGF receptor cells to produce VEGF molecules inside the *i*th cell. The external VEGF cells come from the continuum field *C*(*x, y, t*), which diffuses from *x* = *L*. Let **x**_*i*_ be the pixel of the *i*th cell that is closer to the hypoxic region. The number of external VEGF molecules in that pixel is *C*(**x**_*i*_, *t*) multiplied by the conversion factor *χ_V_* = *N*_*A*_*L*^2^/[(*N* − 1)^2^*M_V_*], where *M_V_* is the molecular weight of the VEGF molecules and *N_A_* is the Avogadro number. We have used *χ_V_* = 1, which is representative of VEGF molecules with a large molecular weight. In the numerical simulation, *C* is known in the grid points and its value at a pixel should be the average value of the four grid points of the pixel. Since these values are quite similar, we adopt the value of *C* at the bottom left grid point of the pixel **x**_*i*_ as *C*(**x**_*i*_, *t*).

The shifted, excitatory and inhibitory Hill functions appearing in Eqs. (14)-(19) are:

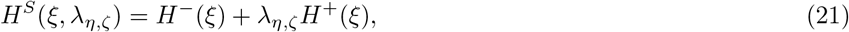

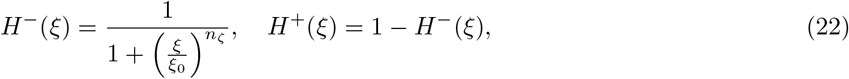

where *H^S^* is excitatory for *λ_η,ζ_* > 1 and inhibitory for *λ_η,ζ_* ≤ 1. In Eqs. (21)-(22), *ξ* = *V, I*, *η* = *I, V, D, J*, and *ζ* = *N, D, J*, *V_R_*, *F* (the subscript *F* refers to Fringe, cf. [47]). The dimensionless parameters *n_ζ_* and *λ_η,ζ_* appearing in the Hill functions are listed in Table 3.

**Table 3.** Dimensionless parameters appearing in the Hill functions. *I*_0_ and *V*_0_ are activation numbers of NICD and VEGF molecules, respectively, and *χ_V_* is the conversion factor.

| Parameter | $\lambda_{I,N}, \lambda_{V,D}, \lambda_{I,J}$ | $\lambda_{I,D}, \lambda_{I,V_R}$ | $\lambda_{D,F}$ | $\lambda_{J,F}$ | $n_N, n_D, n_V, n_{V_R}$ | $n_J$ | $n_F$ | $I_0, V_0$ | $\chi_V$ |
| --- | --- | --- | --- | --- | --- | --- | --- | --- | --- |
| Value | 2.0 | 0.0 | 3.0 | 0.3 | 2.0 | 5.0 | 1.0 | 200 | 1.0 |

### Cell types, proliferation, branching and anastomosis

#### Cell types

In the model, ECs may be on a tip, hybrid or stalk cell phenotype. In nature, tip cells are characterized by having high levels of Delta-4, VEGFR2, and active VEGF signaling (i.e., high levels of VEGF internalization). They develop filopodia and migrate along the VEGF-A gradient, leading the formation of new branches. Delta-4 proteins at tip cell membranes inhibit the neighboring cells (due to lateral inhibition) to adopt a tip phenotype, thereby forcing them to become stalk cells (with low Delta-4, VEGFR2 and internalized VEGF).

Likewise, in our model, tip cells are distinguished by the number of VEGF molecules they possess. Therefore, a cell that has *V* larger than all its neighbors and *V* > 0.5 max_*i*_*V*_*i*_(*t*) will acquire the tip cell phenotype and be very motile. To simulate this, tip cells are able to follow the mechanical and chemical cues on the environment, having *ρ*_chem_ ≠ 0 and *ρ*_durot_ ≠ 0. On the other hand, stalk cells are not motile and they have *ρ*_chem_ = *ρ*_durot_ = 0 in the model (except when they undergo proliferation, see below). Stalk cells, by virtue of the lateral induction, characteristic of Notch-Jagged signaling, are able to induce neighboring cells to adopt a stalk cell phenotype, by promoting a decrease of internal VEGF in them.

In our model we track the cells belonging to each growing vessel. A new sprouting vessel can be formed when a stalk cell acquires the tip phenotype. This cell can then become the leading cell of a new vessel that branches out from out the old one. This is illustrated in Fig. 1. If the levels of VEGF inside the tip cells that lead an active growing branch drop to values in the interval 0.2 max_*i*_*V*_*i*_(*t*) < *V* < 0.5 max_*i*_*V*_*i*_(*t*), these cells will be in the hybrid phenotype. In spite of the lower amount of Delta-4, VEGFR2 and VEGF, these cells remain with the tip cell characteristics and are able to lead the sprout. Similarly, stalk cells whose internal VEGF increases to the same range acquire the hybrid phenotype and can lead a sprout. The number of cells in the hybrid phenotype is only appreciable for larger Jagged production rates.

#### Branching

When a stalk cell (which does not border an existing tip cell) acquires the tip cell or the hybrid tip/stalk cell phenotype, this event will lead to the creation of a new active sprouting branch depending on its localization within the existing branch and on its moving direction.

To create a new branch, the boundary of the tip cell must touch the ECM. Moreover, let 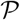 be the set of 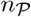 ECM pixels that have boundary with the branching tip cell. For each pixel 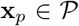, let the *strain vector* be **v**_*p*_ = *ϵ_j_* **v**_*j*_, where *ϵ_j_* is the largest eigenstrain at pixel **x**_*p*_ and **v**_*j*_ is the corresponding unit eigenvector, as defined after Eq. (3). The average modulus and argument for the branching cell *i* are

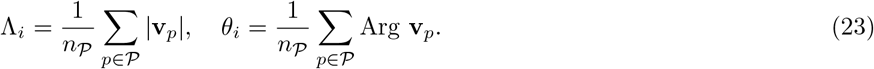

Let us also assume that the gradient of the chemotactic factor *C* forms an angle Θ with the *x*-axis. The new tip cell will branch out, creating a new vessel, if the direction given by *θ_i_* points in the direction of the ECM and if −*π*/2 < *θ_i_* − Θ < *π*/2. For other values of *θ_i_*, the tip cell does not leaves the parent vessel, since the chemotactic term of Eq. (4) opposes branching. In those cases, the direction given by *θ_i_* points to another cell, not to the ECM. To facilitate branching computationally, we directly exchange the new tip cell with this neighboring cell (see Figs. 1(a) and 1(b)). These exchanges may continue in successive MCTS until the new tip cell reaches a boundary of the blood vessel for which the direction given by *θ_i_* points to the ECM, as shown in Fig. 1. These cell exchanges in 2D mimic the climbing motion of the new tip cell over the parent vessel in a 3D geometry without merging with it.

We set the branching process to take at least 400 MCTS (incubation time). During this time we implement a persistent motion of the new tip cell in the direction marked by the angle *θ_i_*. During this incubation period, and to provide a good separation from the parent vessel we permit the tip cell to proliferate once (see Figs. 1(e) and 1(f), and see below). After this time, the dynamics of the branching vessel follows the same rules as that of any other actively sprouting vessel.

#### Cell proliferation

Endothelial cell proliferation in sprouting angiogenesis is regulated by both mechanical tension and VEGF concentration. In sprouting angiogenesis the tip cell creates tension in the cells that follow its lead. On those first stalk cells, this tension produces strain that triggers cell proliferation, if VEGF concentration is high enough [35]. Therefore, in our model, for each active sprouting vessel, one of the stalk cells that is in contact with a tip cell is randomly chosen to undergo proliferation. Only one cell per sprout proliferates, and it takes an average of 20 MCTS to do so. Note that while it takes an average 20 MCTS (0.88 hours) for a cell to proliferate, the video frames in the Supplementary Material are taken every 50 MCTS, which gives the wrong impression of cell proliferation faster than experimentally observed. In fact, the mean cell proliferation time of 0.88 hours in the simulations is within the experimentally observed span between 0.3 and 1.2 hours [83]. Tip cells in the model cannot proliferate, except only once when they start a new branch.

Once a stalk cell attached to a tip cell has been randomly selected as a proliferating cell, its target area in the CPM is set to become twice the size, whereas its target perimeter is set to a value three times that of non-proliferating cells. This cell will then grow in successive MCTS until it reaches this large target area. Then the cell proliferates if the following three conditions hold: (i) *C*(**x**_*i*_, *t*) > *ψ*_*p*_ (external VEGF surpasses a threshold), (ii) the cell belongs to an active blood vessel with cell proliferation, and (iii) the cell is not surrounded completely by other cells. Failure to meet one of these conditions precludes proliferation. If the three conditions are met, we use the unsupervised machine learning algorithm *K-means clustering* to split the cell. This algorithm calculates the Euclidean distance of each pixel in the cell to the centroid of two groups of pixels and corrects the centroids until the two pixel groups are balanced. These two groups comprise the new cells. Provided the daughter cells share boundary with the tip cell, one of them is randomly chosen to retain the ability to proliferate but the other cell does not proliferate. If the daughter cells do not share boundary with the tip cell, they both become non-proliferating and a different cell that shares boundary with the tip cell is randomly chosen to become a proliferating cell. Non-proliferating stalk cells have *ρ*_chem_ = *ρ*_durot_ = 0, whereas proliferating stalk cells are affected by chemo and durotaxis and their *ρ*_chem_ and *ρ*_durot_ values are as in Table 1. Setting *ρ*_chem_ = *ρ*_durot_ = 0 for all proliferating stalk cells impedes branching of new sprouting vessels that are unable to leave the neighborhoods of their parent vessels.

#### Anastomosis

When an active sprouting blood vessel merges with another active sprouting vessel, i.e. during anastomosis, one of them becomes inactive. If the collision occurs between tip cells of two different vessels, one vessel is randomly chosen to become inactive. If one tip cell merges with a stalk cell of a different active sprouting vessel, the vessel to which the tip cell belongs becomes inactive. The cells of an inactive vessel do not proliferate or branch, although they continue to undergo Notch signaling dynamics.

#### Cell motility

In experiments [**?**, 82, 83], observed endothelial cell motility is much higher than in our simulations. Real cells move in 3D and can climb past each other but our simulations are 2D. 2D geometry necessarily and artificially limits cell motility, as cells cannot interpenetrate each other. We have overcome this limitation in the important case depicted in Fig. 1, when a cell that has just acquired the tip cell phenotype finds itself constrained by a stalk cell that blocks its way to reach the boundary of the vessel with the ECM and branch out. In this case, we exchange neighboring cells, which is a way to mimic the 3D effect of the tip cell moving past the blocking stalk cell. While we could modify the 2D model postulating stalk cell motility and exchanges among stalk cells to mimic their observed 3D cell motility, this would complicate our 2D model. Note that the current model already reproduces the observed growth of blood vessels, without having to consider stalk cell motility and stalk cell exchange. More realistic cell motility would be addressed in future 3D versions of the current model.

## Results and discussion

The simulations of our model were implemented on Graphics Processing Units (GPU) using C-CUDA (CUDA: Compute Unified Device Architecture created by NVIDIA Corporation). This software contains source code provided by NVIDIA Corporation. The visualization of the results uses Matlab. We have elaborated our own simulation code, which is based on that by van Oers *et al.* [66] (implemented in the programming language C with Matlab visualization), K-means CUDA algorithm [85], standard algorithms (Euler, finite differences and finite elements) to solve ordinary and partial differential equations and CUDA libraries that are specified in the Supplementary Material. The flow diagram of the model is presented in Fig. 2, which encompasses the different computational modules for branching, cell proliferation, VEGF concentration, cell signaling processes, mechanics, CPM and anastomosis, cf the Supplementary Material. Due to the complexity of the model, parallel computing using C-CUDA allows the reduction of the computational times as much as possible. The amount of processes that can be calculated at the same time (over pixels, cells, vessels…) make this problem manageable. Furthermore, the implementation of our own code allow us to control times, features, parallel processes and the addition or changes of modules. The computation time of each reported simulation in a computer with Intel(R) Core(TM) i7-7700K CPU @4.20 GHz processor, 64.0 GB RAM and NVIDIA GeForce GTX 1080 graphics card is about 4 hours. We have run our simulation model for a simple slab geometry and different conditions. A primary vessel is supposed to be along the *y* axis. The initial VEGF concentration *C*(**x**, 0) is independent of *y* and decays linearly in *x* from *x* = *L* to *x* = 0. Thus, chemotaxis pushes tip cells towards the vertical line at *x* = *L*.

**Figure 2.**
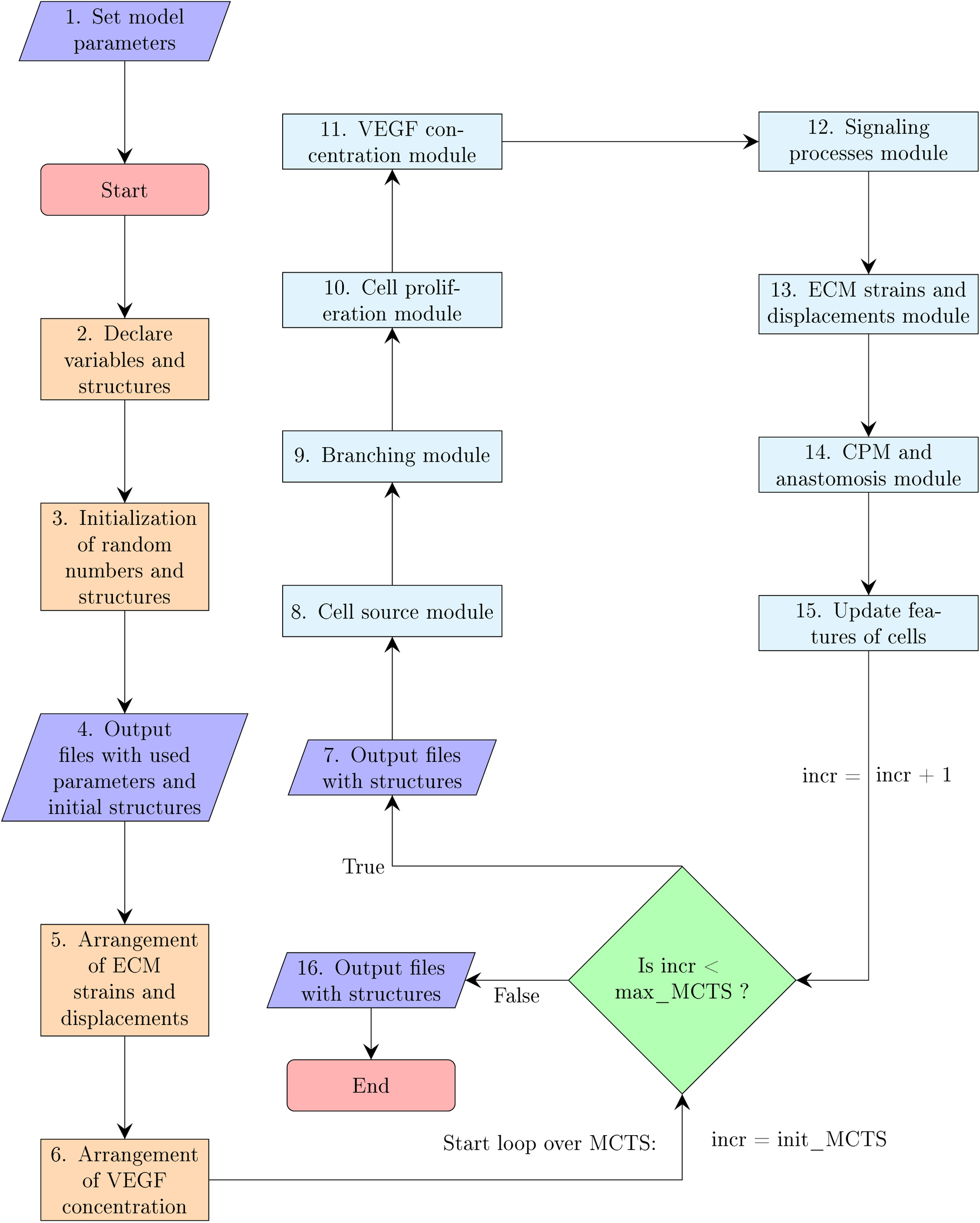
Flow diagram of the simulation model.

### Cellular mechanics and anastomosis

It is clear that without the deformation of ECM induced by cells tractions and the strain vectors, there will be no branching direction for new tip cells to exit from a given sprout. Thus, cellular mechanics is crucial for branching. We have also found that cellular mechanics significantly controls anastomosis. The arrows in Fig. 3 are directed along the strain vector (eigenvector corresponding to the largest eigenstrain and having length equal to that eigenstrain). According to Eq. (3), the arrows indicate the likeliest direction in which ECs will move. The snapshots depicted in Fig. 3 show examples of successful and frustrated anastomosis and branching of advancing blood vessels. A tip cell leads successful branching from the blood vessel at the bottom of Fig. 3(a), as shown by Panels (b) and (c). Meanwhile, two blood vessels that sprout from the blood vessel at the top of Fig. 3(a) successfully anastomose as shown in Fig. 3(c). Notice that the strain vectors show the path of the approaching vessels until they anastomose. However, the branches arising from the two lowest vessels in Fig. 3(a) do not anastomose. They approach each other in Fig. 3(c) but the strain vectors pull them away from each other and anastomosis is frustrated, as shown in Figs. 3(d) and 3(e) [66, 86, 87].

**Figure 3.**
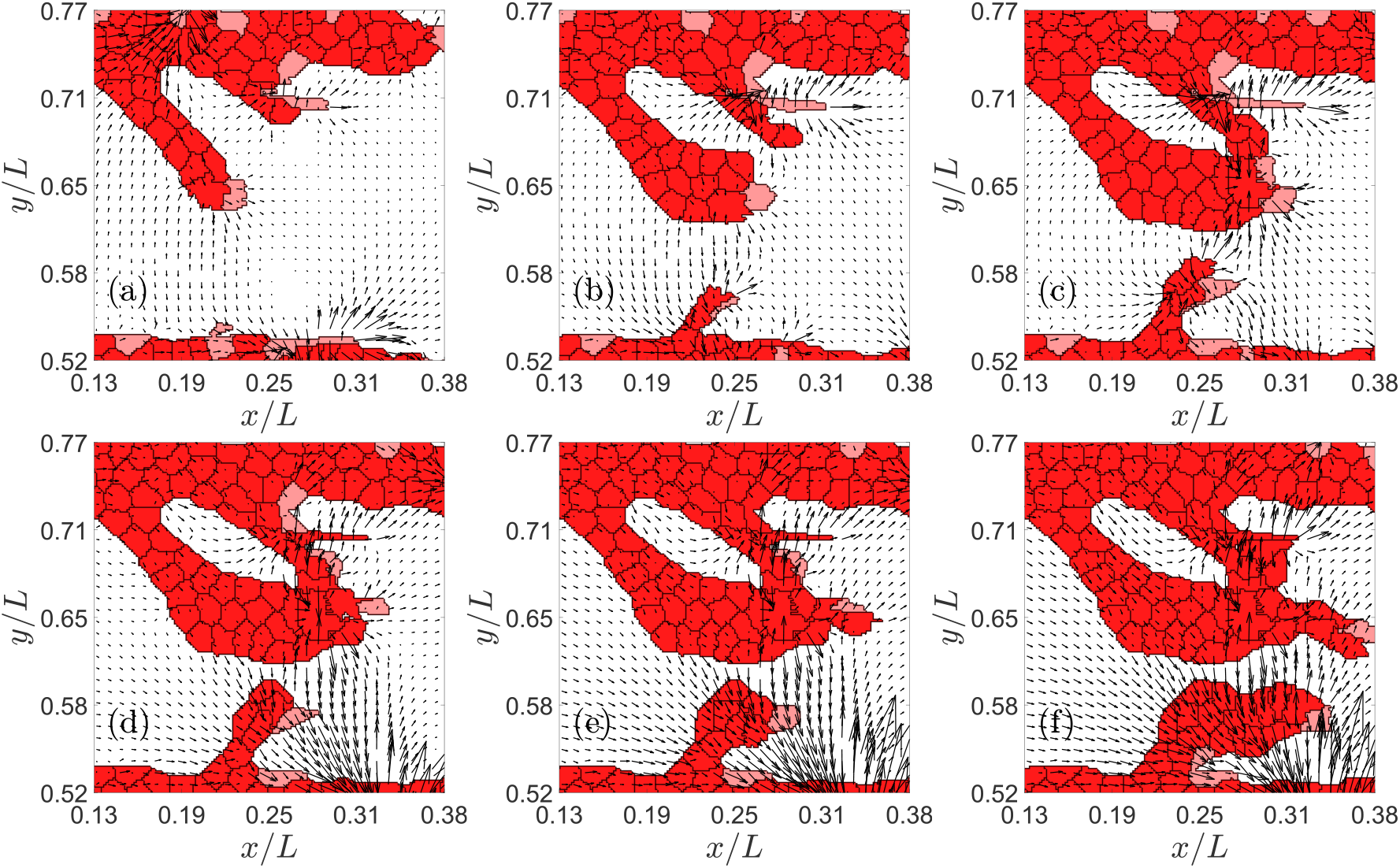
Example of successful and frustrated anastomosis. Times in MCTS are: (a) 751, (b) 851, (c) 951, (d) 1051, (e) 1101, (f) 1201. Tip cells are pink.

Tip cells have higher levels of VEGF and their motion follows stiffness, chemical and adhesion gradients, as expected from the model. In successful anastomosis, one tip cell is directed by the strain vector to one actively sprouting vessel. When it makes contact, it fuses with that vessel. After that, the VEGF in the tip cell decreases and it becomes a stalk cell.

### Jagged–Delta dynamics and sprouting

Jagged and Delta dynamics determine sprouting [47, 48]. Studies of Notch signaling in one cell driven by external Jagged and Delta molecules show that the phenotype of a tip cell changes to hybrid tip/stalk and then to stalk cell as the external Delta concentration surpasses successive thresholds (cf. Fig. 3 of Ref. [47]). The thresholds depend on the Jagged production rate. Lateral induction works similarly for one cell driven by external Jagged molecules: tip cells change to hybrid tip/stalk and stalk phenotypes as the external Jagged concentration surpasses successive Delta-dependent thresholds [47, 48]. Simulations of our model illustrate the effects of J-N and D-N signaling combined with chemo-, hapto- and durotaxis. Figure 4 shows that increasing the Jagged production rate *r_J_* yields smaller branching blood vessels, thereby decreasing the irrigation of the hypoxic region. Furthermore, sprouting is accelerated as the Jagged production augments: thinner and less efficient sprouts are formed faster as *r_J_* increases. Stalk cells proliferate on advancing sprouts. Thus, increasing the number of tip cells leading sprouts results in increasing cell proliferation and a more rapid sprout advance. This behavior agrees with the sketch in Fig. 5A of Ref. [47], which indicates that pathological angiogenesis is obtained when there is an excess Jagged production. The sprouts in physiological angiogenesis are thicker and advance more slowly than the more abundant and thinner sprouts in pathological angiogenesis, as shown in Fig. 4.

**Figure 4.**
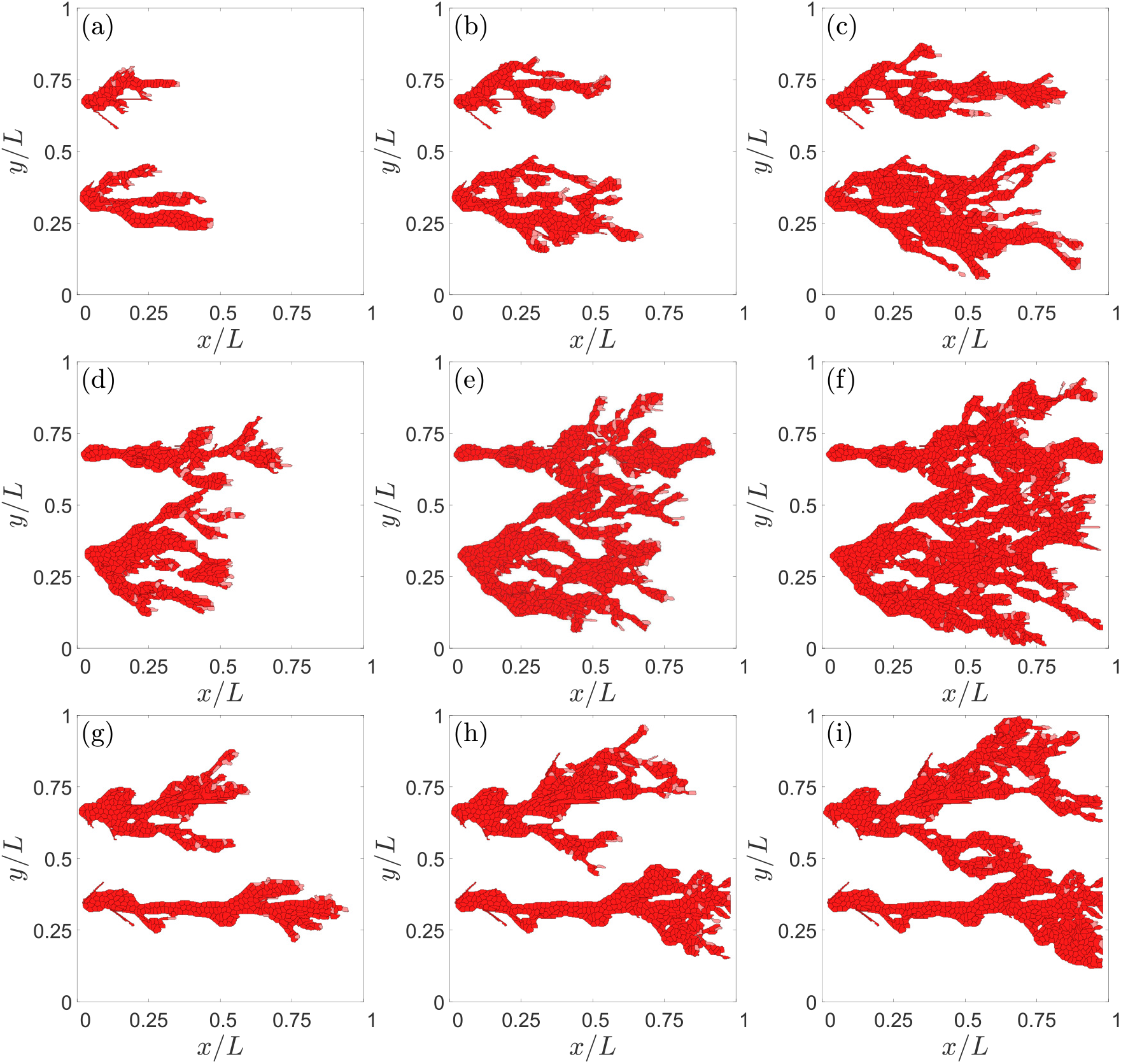
Effect of Jagged production on angiogenesis. For *r_J_* = 500 molec/h and *r_D_* = 1000 molec/h, snapshots at times: (a) 2001 MCTS, (b) 2751 MCTS, (c) 3501 MCTS. For *r_J_* = 2000 molec/h and *r_D_* = 1000 molec/h, snapshots at times: (d) 2001 MCTS, (e) 2751 MCTS, (f) 3501 MCTS. For *r_J_* = 2000 molec/h and *r_D_* = 7500 molec/h, snapshots at times: (g) 2001 MCTS, (h) 2501 MCTS, (i) 3501 MCTS.

**Figure 5.**
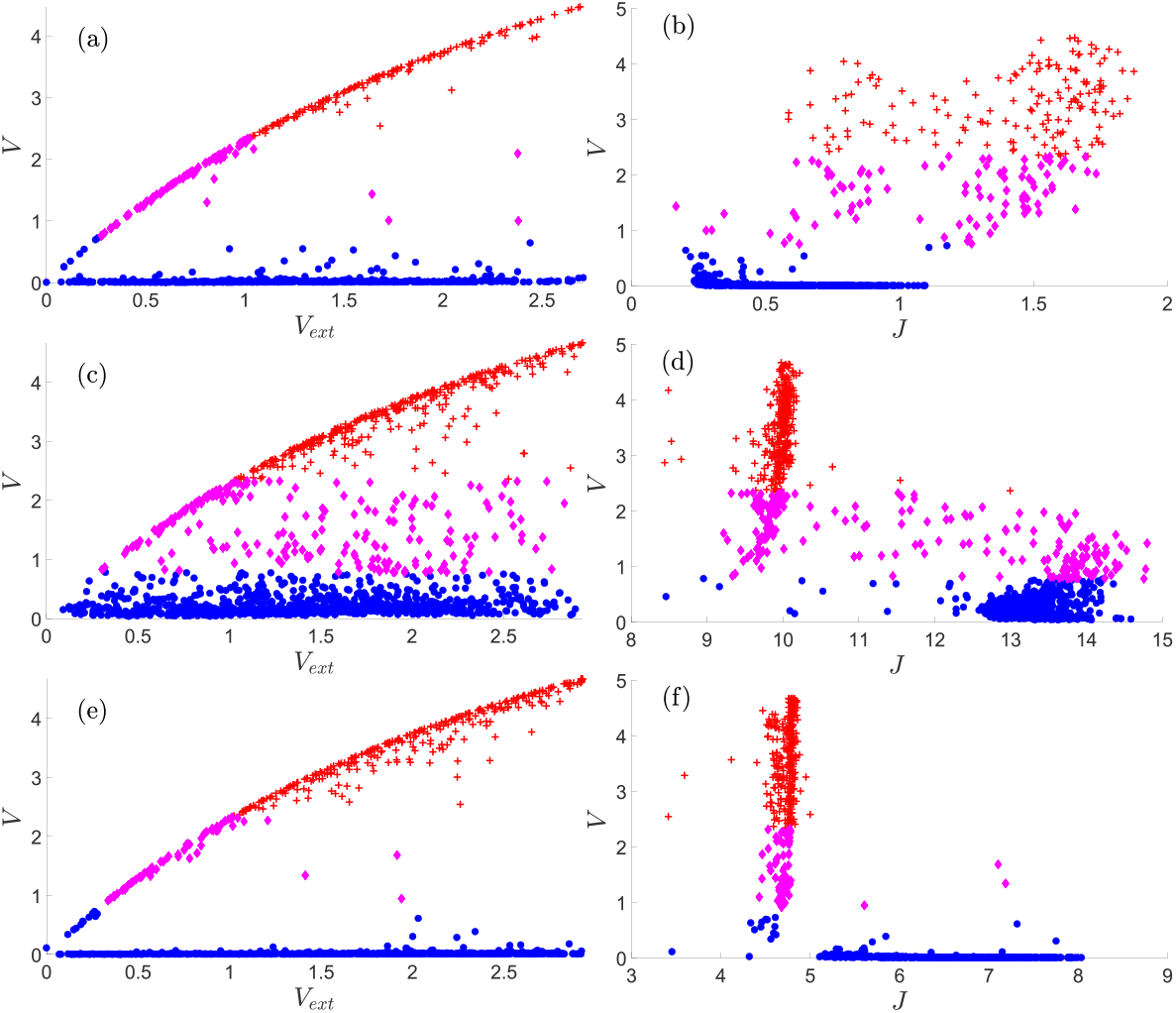
Content of VEGF, *V*, versus *V*_ext_ = *C*, and of *V* versus *J* in the tip, stalk and hybrid tip-stalk cells within the angiogenic network at 3501 MCTS. (a), (b) *r_J_* = 500 molec/h, *r_D_* = 1000 molec/h; (c), (d) *r_J_* = 2000 molec/h, *r_D_* = 1000 molec/h; (e), (f): *r_J_* = 2000 molec/h, *r_D_* = 7500 molec/h. Other parameter values are as indicated in Tables 1-3. Nondimensional units for *V*, *V*_ext_, *J* are as indicated in Table 4. The meaning of symbols is as follows. Red cross (tip cell), magenta rhombus (hybrid tip/stalk cell), blue circle (stalk cell).

The Delta production rate *r_D_* acts opposite to *r_J_*. High and intermediate levels of *r_D_* ensure physiological angiogenesis, whereas the numbers of the hybrid tip/stalk cells increase for low levels of *r_D_*. In more detail, we observe that, for *r_J_* = 500 molec/h and *r_D_* = 1000 molec/h, Figs. 5(a) and 5(b) show a gap between the VEGF of stalk and tip cells: the content of *V* is very low for stalk cells. It increases monotonically with *V*_ext_ and *J* for hybrid-tip and tip cells. Fig. 5(b) also shows that tip cells and hybrid tip-stalk cells have larger *J* than stalk cells. As *r_J_* increases, at *r_J_* = 2000 molec/h, the hybrid-tip cells have proliferated and bridge the gap in *V*, as depicted in Figs. 5(c) and 5(d). Fig. 5(d) indicates that *J* is smaller for the tip cells at large Jagged production rates, which is consistent with lateral induction of stalk phenotype by stalk cells with large *J* values [47]. For large *r_J_*, tip cells have less Jagged (*J* ≈ 10 and *V* > 2) than other cell types (*J* between 10 and 15 and *V* < 2), as shown in Fig. 5(d). Fig. 5(e) and 5(f) show that the Delta production rate *r_D_* acts in opposition to *r_J_*. At *r_D_* = 7500 molec/h, Fig. 5(e), there is again a gap between the VEGF of stalk and tip cells. At this large Jagged production rate, tip cells have lower Jagged than stalk cells, as depicted in Fig. 5(f), which is similar to Fig. 5(d). However, Fig. 5(f) exhibits a gap between the maximum value of *J* for tip cells and the values of *J* for stalk cells, as compared to Fig. 5(d).

**Table 4.** Units for nondimensionalizing the Notch equations (14)-(19).

| Variable | $N_i, D_i, J_i, N_{\text{ext}}, D_{\text{ext}}, J_{\text{ext}}$ | $I_i$ | $V_{Ri}$ | $V_i$ | $V_{\text{ext}}$ | $t$ |
| --- | --- | --- | --- | --- | --- | --- |
| Scale | $\sqrt{r_D/k_C}$ | $(k_T r_D)/(k_C \gamma_S)$ | $r_{V_R}/\gamma$ | $V_0$ | $6V_0$ | $1/\sqrt{k_C r_D}$ |
| Value | $\sqrt{2} \times 10^3$ | $10^2$ | $10^4$ | $2 \times 10^2$ | $12 \times 10^2$ | $\sqrt{2}$ |
| Unit | molec | molec | molec | molec | molec | h |

### Jagged–Delta dynamics and anastomosis

What is the effect of modifying J-N and D-N signaling on angiogenesis? Figs. 4 and 6 show the effects of lateral inhibition by D-N signaling versus lateral induction by J-N signaling. Increasing the Jagged production rate produces more hybrid tip/stalk cells and more sprouts, as shown by Figs. 4 and, for the higher content of hybrid tip/stalk cells, by Figs. 5(c) and 5(d). However, for a high Jagged production rate, increasing the Delta production rate favors lateral inhibition by tip cells, which eventually decreases the number of new sprouts, makes anastomosis less frequent, as illustrated by Figs. 4(g), 4(h), 4(i), and 6.

**Figure 6.**
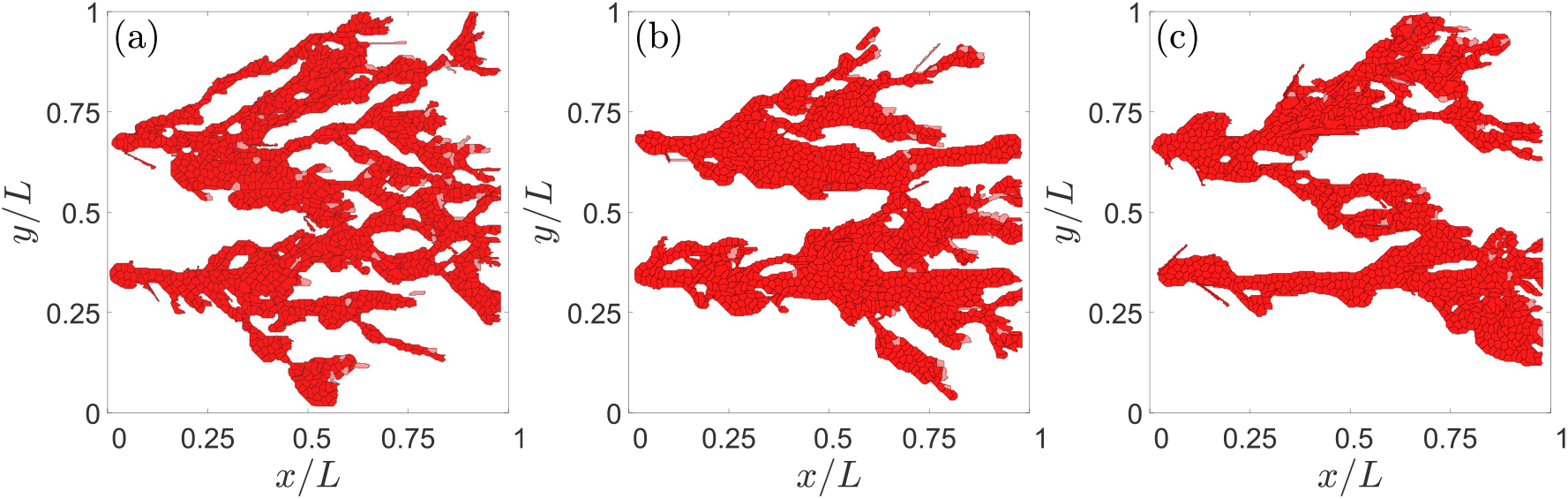
Effect of the Delta production rate on angiogenesis with a high Jagged production rate of *r_J_* = 2000 molec/h at 3501 MCTS. (a) *r_D_* = 3000 molec/h, (b) *r_D_* = 6000 molec/h, and (c) *r_D_* = 7500 molec/h. Lateral inhibition due to more activated D-N signaling decreases the number of hybrid tip/stalk cells and branching.

Fig. 7 shows the concentrations of *N*, *V*, *J* and *D* for a developed angiogenic network for several values of the Jagged-1 and Delta-4 production rates. We observe that tip cells have large values of *V* and *D* for both normal (*r_J_* = 500 molec/h) and high (*r_J_* = 2000 molec/h) Jagged production rates, cf. Figs. 7(g)-(i) and 7(m)-(o). These figures highlight the role of lateral inhibition on stalk cells that are neighbors of tip cells. For large *r_J_*, Figs. 7(k) and 7(l) show that stalk cells clearly have larger values of *J*, thereby illustrating the more important role of lateral induction. For large *r_J_* and moderate *r_D_*, Fig. 7(h) exhibits a larger number of cells with intermediate values of their internal VEGF, which shows the abundance of the hybrid tip/stalk cell phenotype. This is not the case for lower Jagged production rate as shown by the VEGF content in Figs. 7(g) and, for higher Delta production rate, in Fig. 7(i). As explained before and as shown by comparing Figs. 5(b) to 5(d) and 5(f), stalk cells have a smaller value of *J* than tip or hybrid tip/stalk cells at smaller Jagged production rates. In these cases, lateral inhibition by D-N signaling is more important. Figs. 7(m), 7(n) and 7(o) show that the *D* level of stalk cells is much reduced as compared with that of neighboring tip cells. Increasing the production rate of Delta-4 restores the morphology of the advancing normal vasculature to angiogenesis with high Jagged production rate, as shown by a comparison of Fig. 7(n) to Figs. 7(o) and 7(m).

**Figure 7.**
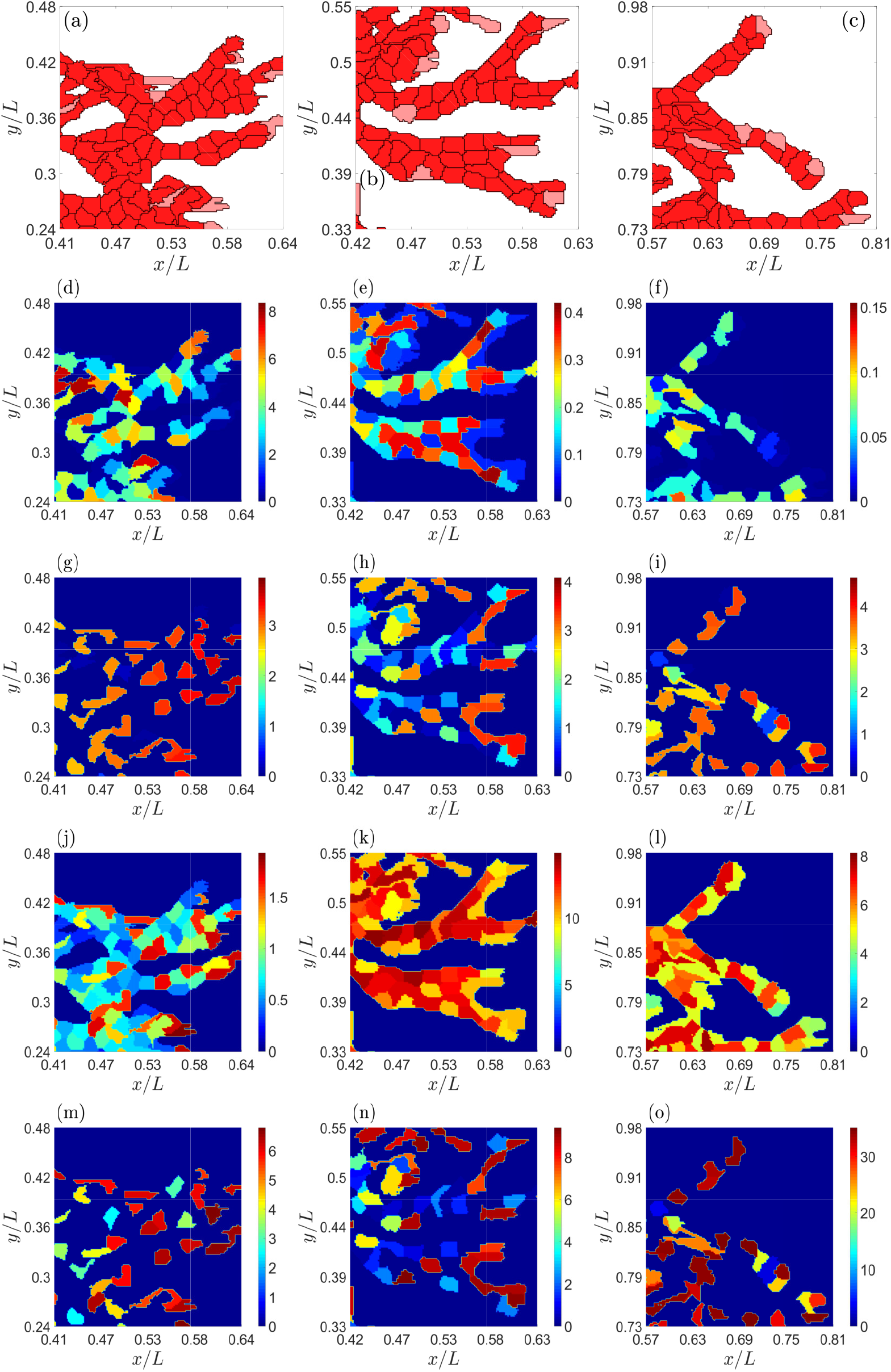
Effect of the Jagged and Delta production rates on angiogenesis at a time of 2901 MCTS. (a)-(c) Snapshots of networks, (d)-(f) Notch concentration, (g)-(i) VEGF concentration, (j)-(l) Jagged-1 concentration, (m)-(o) Delta-4 concentration. Data: (a),(d),(g),(j),(m) *r_J_* = 500 molec/h, *r_D_* = 1000 molec/h, (b),(e),(h),(k),(n): *r_J_* = 2000 molec/h, *r_D_* = 1000 molec/h, (c),(f),(i),(l),(o): *r_J_* = 2000 molec/h, *r_D_* = 7500 molec/h. Nondimensional units for protein concentrations are as in Table 4.

Figure 8 further shows the effect of varying the production rates of Jagged and Delta on the advance and morphology of the vascular plexus. With respect to the simulations in Figs. 1 and 3 for standard values of *r_J_* and *r_D_*, increasing the production of Jagged, as shown in Fig. 4, produces more tip cells that run faster, cf. Figs. 8(a) and 8(b). Thus, lateral induction mediated by Jagged accelerates the advance of vasculature and increases the number of blood vessels by creating more hybrid tip/stalk cells, as explained before in relation to Figs. 4 and 5. If we keep constant *r_J_* and increase the Delta production rate, lateral inhibition by tip cells becomes stronger, cf. Fig. 4. Then the number of tip cells decreases whereas the vasculature advances only slightly faster because angiogenesis and anastomosis diminish compared with the case of smaller *r_D_*, cf. Fig. 8(c) and 8(d).

**Figure 8.**
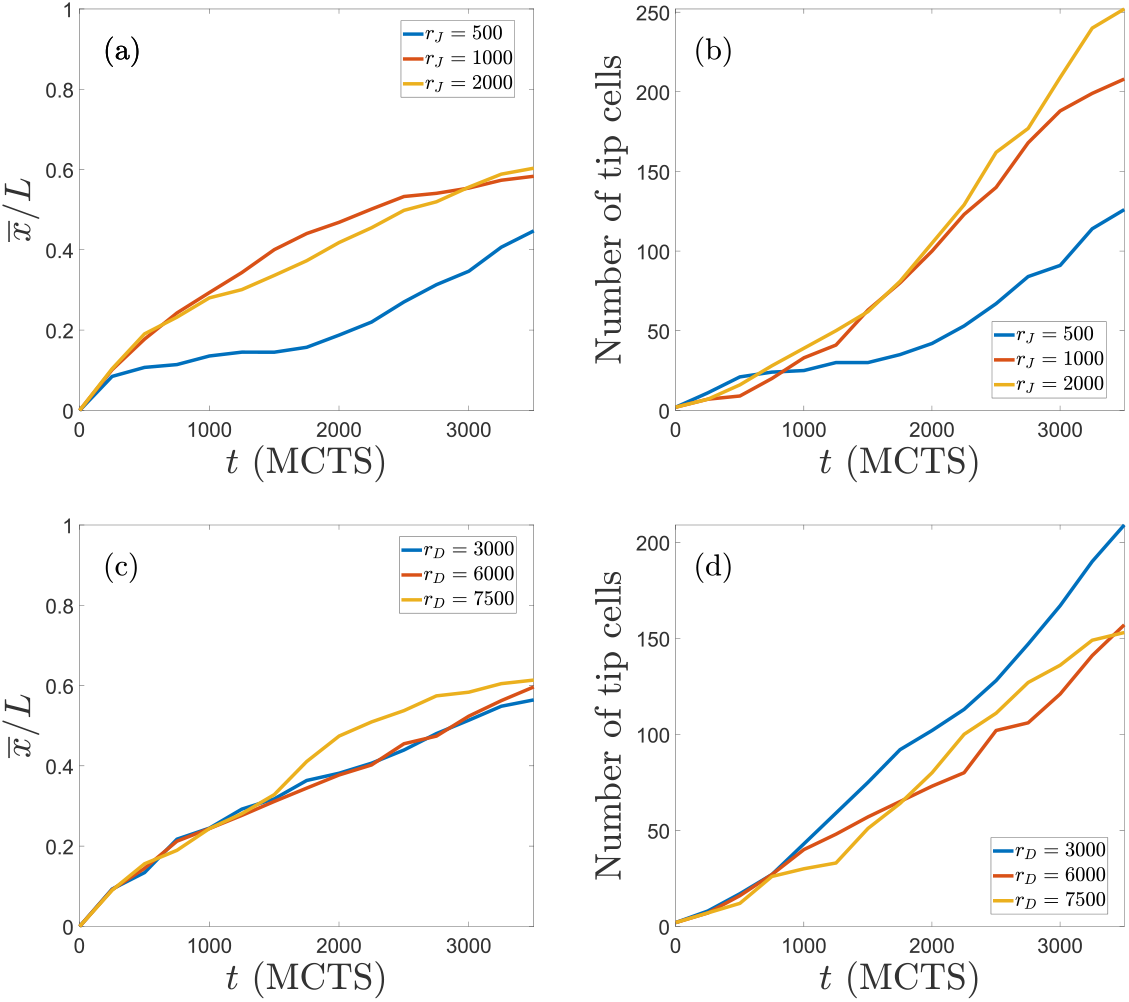
(a) Average abscissa (position on *x* axis) of the tip cells as a function of time, and (b) number of tip cells versus time for *r_D_* = 1000 molec/h and *r_J_* = 500, 1000 and 2000 molec/h. Increasing Jagged production rate yields more tip cells that advance faster. (c) Average position of tip cells versus time, and (d) number of tip cells versus time, for *r_J_* = 2000 molec/h and *r_D_* = 3000, 6000 and 7500 molec/h. Increasing Delta production rate makes tip cells to advance slightly more but it diminishes the number of tip cells. The effect of *r_D_* on the number of tip cells is opposite to that of *r_J_* in Panels (a) and (b).

### Sensitivity

The sensitivity of the results to the particular parameter set chosen is studied by varying one parameter at a time. In previous paragraphs, we have analyzed the effect of varying Potts parameters on the simulations of the model. They affect the relative importance of mechanical and chemical cues as described, and their effects are consistent with previous works on chemotaxis [76] and durotaxis [66]. Here we discuss the sensitivity of simulation results to changes in the parameters controlling cellular signaling. To this end, we have carried out 6 simulations for each of the production rates mentioned above and taken the averages of these realizations. Figs. 9(a) and 9(b) display, as a function of time, the number of angiogenic sprouts and the percentage of pixels of the hypoxic region at *x* = *L* that are occupied by them, respectively. The number of sprouts and the occupation fraction *φ* should be contrasted with Figs. 4 to 6. For fixed *r_D_* = 1000 molec/h, increasing *r_J_* produces thinner and more numerous pathological sprouts that arrive faster to *x* = *L*. Increasing *r_D_* at a higher *r_J_* decreases the proliferation of sprouts and the fraction of pixels occupied by them at the hypoxic region. However, the sprouts move faster towards the hypoxic region, which keeps having a higher occupation fraction *ϕ* than in the case of physiological angiogenesis with lower *r_J_*. Increasing Delta production decreases the number of sprouts (and thickens them), as corroborated by Ubezio *et al*’s experiments [45]. Fig. 10 depicts how the percentages of tip and stalk cells in moving sprouts evolve in time for the data of Fig. 9. In all cases, the percentages stabilize to the same low values of tip cells and high values of stalk cells after 2000 MCTS (time it takes the first sprouts to arrive at the hypoxic region). For shorter times, the influence of production rates on the relative number of tip/stalk cells is evident: higher *r_J_* lowers the percentage of tip cells, whereas the influence of an increment of *r_D_* on the percentage of tip/stalk cells is less clear. These data need to be contrasted with those of Figs. 4 to 6 to achieve a clearer picture of the morphology and thickness of the angiogenic network.

**Figure 9.**
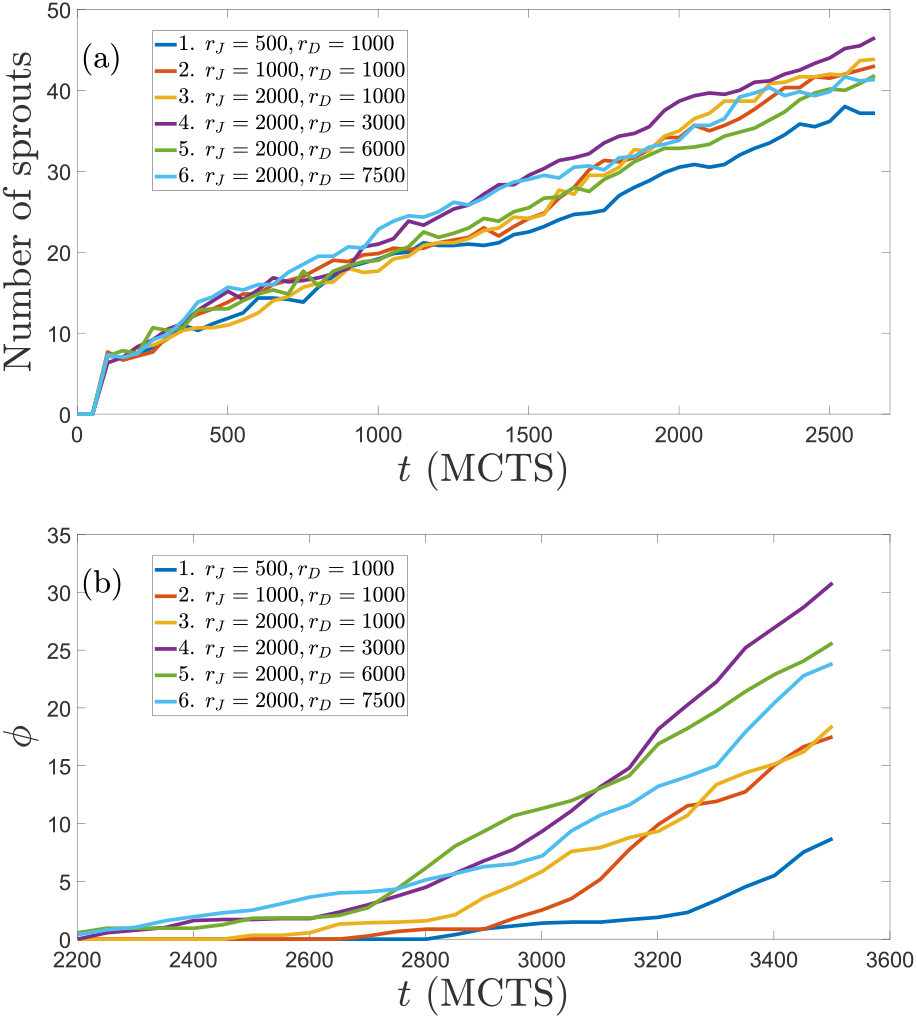
(a) Number of angiogenic sprouts versus time, and (b) percentage of pixels *ϕ* at *x* = *L* (the hypoxic region) that are occupied by vessel sprouts versus time, for the indicated Jagged and Delta production rates. Data correspond to averages over 6 realizations of the stochastic process.

**Figure 10.**
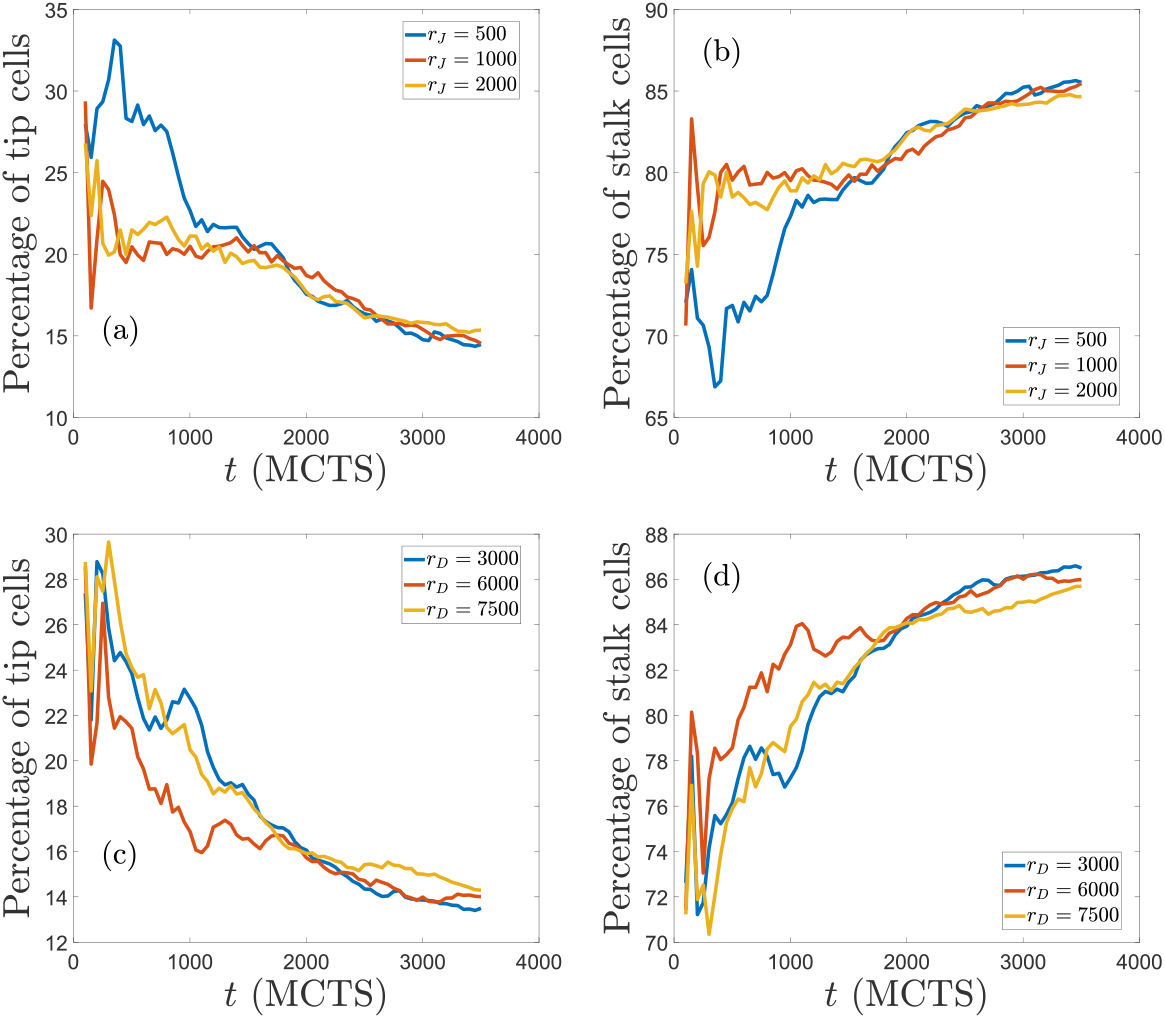
Percentage of tip and stalk cells versus time for the simulations displayed in Fig. 9. Production rates are *r_D_* = 1000 molec/h in panels (a) and (b), and *r_J_* = 2000 molec/h in panels (c) and (d).

## Conclusion

The mathematical model of angiogenesis presented here illustrates the relative importance of mechanical, chemical and cellular cues when they are all considered simultaneously. Given a proliferation rate and a VEGF gradient on a homogeneous extracellular matrix, competing J-N and D-N dynamics determine the influence of lateral inhibition and lateral induction on tip cell selection, branching, anastomosis and speed of angiogenesis. Anastomosis is driven by chemotaxis. Cellular motion is informed by haptotaxis and durotaxis. However, anastomosis may be favored or impeded depending on the mechanical configuration of strain vectors in the ECM near tip cells. Notch signaling determines tip cell selection and vessel branching. Lateral induction by stalk cells and lateral inhibition by tip cells are informed by competing Jagged-Notch and Delta-Notch dynamics in manners that depend quantitatively on the Delta and Jagged production rates. In particular, the numerical simulations of our model predict the following effects of the production rates. Increasing the production rate of Jagged favors lateral induction of stalk cells, which yields more hybrid tip/stalk cells and a thinner vasculature that advances faster. On the other hand and as observed in experiments [45], increasing the production rate of Delta lowers the number of tip cells by lateral inhibition of stalk cells. Then there are less sprouts and anastomosis is less frequent while the advance of the vascular plexus is only slightly faster. Our numerical simulations illustrate the regulating role of Notch-Jagged-Delta signaling in the velocity and morphology of angiogenic vasculature. An imbalance of the Jagged production, so that there is more Jagged and increased lateral induction of stalk cells, results in anomalous thinner sprouts and faster angiogenesis. This may be corrected by increasing the Delta-4 production rate, which boosts lateral inhibition of tip on stalk cells, diminishes the number of tips and slows down somewhat angiogenesis.

To allow for quantitative comparisons with experiments, e.g., [45], our 2D model of early stage angiogenesis needs to be extended in several directions to be made more realistic and to account for later stages of angiogenesis. The extension of the model to three dimensional configurations is straightforward although it requires more computing power. While we have studied relatively short distances between the primary vessel and the target hypoxic region, we need to consider larger systems to be able to do statistical studies of vessel numbers and their width. To move toward later stages of the formation of an advancing vascular plexus, we need to add lumen formation [24] and blood circulation to the model [31]. These processes will allow us to tackle the concurrent sprouting and anastomosis on the front of the advancing vascular plexus and the pruning of poorly perfused sprouts on its back [31, 32].

## Supporting information

description of numerical code

## Acknowledgements

This work resulted from a collaboration that began at the Workshop on Modeling Biological Phenomena from Nano to Macro Scales, held in 2018 at the Fields Institute in Toronto, Canada. We thank Prof. I. Hambleton, director of the Fields Institute, for the invitation and support during the Workshop, and also Prof. A. Carpio, the organizer of the Workshop. LLB thanks Russel Caflisch for hospitality during a sabbatical stay at the Courant Institute of Mathematical Sciences, New York University.

## Supporting Information

**S**1 Text containing information on the numerical code. S2 Supporting movies with a README text file.

## Notes

#### Summary of Updates

3 new figures, revised text.

