## Supplementary material for "Notch signaling and taxis mechanims regulate early stage angiogenesis: A mathematical and computational model": description of numerical code

### Documentation of the simulation code

Rocío Vega[1], Manuel Carretero[1], Rui D.M. Travasso[2], Luis L. Bonilla[1,3]

**1 - G. Millán Institute for Fluid Dynamics, Nanoscience & Industrial Mathematics, and Department of Materials Science & Engineering and Chemical Engineering, Universidad Carlos III de Madrid, Avenida de la Universidad 30, 28911 Leganés, Spain**

**2 - CFisUC, Department of Physics, University of Coimbra, R. Larga, 3004-516 Coimbra, Portugal**

**3 - Courant Institute of Mathematical Sciences, New York University, 251 Mercer St, New York, N.Y. 10012, USA**

The model of our manuscript was implemented on Graphics Processing Unit (GPU) using C-CUDA (CUDA: Compute Unified Device Architecture created by NVIDIA Corporation). The visualization is performed using Matlab. This software contains source code provided by NVIDIA Corporation. Our simulation code is based on the simulation code given by van Oers *et al.* [4] (implemented in the programming language C with the visualization in Matlab), K-means CUDA algorithm [3] and some CUDA libraries that will be specified later. Due to the complexity of the model, the parallel computing using C-CUDA allow us to reduce the computational times as much as possible. The amount of processes that can be calculated at the same time (over pixels, cells, vessels...) make this problem manageable. Furthermore, the implementation of our own code allow us to control times, features, parallel processes and the addition or changes of modules.

We show below a flowchart of our simulation code. Each part of the flowchart will be described in detail afterwards.

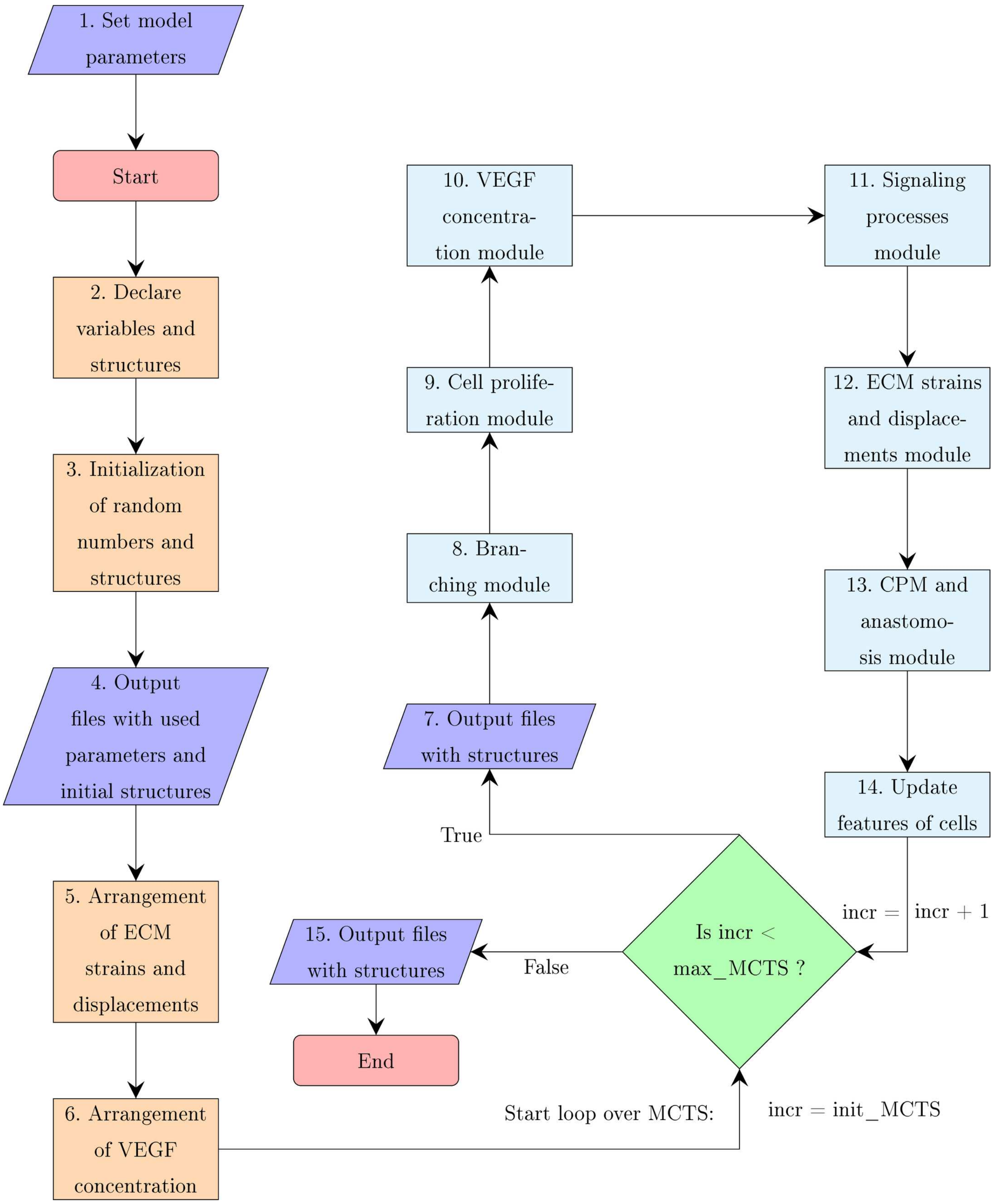


Figure 1: Flowchart.

1. Set model parameters & 2. Declare variables and structures

A description of the files, structures and matrices that make up the simulation code helps to understand the first two parts of the flowchart.

This simulation code consists of the following source (.cpp, .cu,.cc) and header (.h,.cuh) files:

- def.h: defines model parameter values.
- struct.h: defines structures used in the code.
- functions.h : declares all the functions used in the code executed in Central Processing Unit (CPU) (and shows in which .cpp,.cu or .cc file are defined).
- functions_CUDA.cuh: declares all the functions used in the code executed in GPU (and shows in which .cpp,.cu or .cc file are defined).
- cpmfem.cpp: contains the main() function, which calls all other functions of the model.
- init.cpp: initializes pixels, nodes, cells and vessels structures at zero.
- init.cu: sets the initial distribution of cells in pixels, nodes, cells and vessels structures, impose external forces and constrains. This file also contains functions that copy from CPU to GPU or vice versa and then call functions in write.cpp to save output.
- read.cpp: loads input, for instance cell positions.
- write.cpp: saves output, such as cell positions and strains.
- cellmoves.cu : cellular Potts movement and anastomosis.
- cellforces.cu: calculates cell traction forces.
- FE_local.cpp: element stiffness matrices (and using them to calculate stresses and strains).
- FE_assembly.cpp: assembles element stiffness matrices into global stiffness matrix.
- Fe_nodes2dofs.cu: some bookkeeping between the set of elasticity equations and nodal forces and displacements.
- branching.cu: performs the branching of a blood vessel.

- chemotaxis.cu: contains the Preconditioned Conjugate Gradient (PCG) method. This file recalculates the VEGF concentration.
- notchsignaling.cu: calculates number of proteins related to the signaling processes for each cell.
- proliferation.cu: splits in two the cells that meet some conditions.
- fp_abstraction.h, kmcuda.cc, kmcuda.h, metric_abstraction.h, private.h, traspose.cu, tricks.cuh, wrappers.h : files related to K-means algorithm.
- mt.cpp: contains the Mersenne twister algorithm for generation of pseudorandom numbers.

The structures and arrays used to store data of the model are the following:

- VOX: structure with data related to pixels. Arrays contain $(M-1)^2$ elements, where $M$ is the number of nodes on a side of the square domain.
  - ctag: (int array) contains the id of occupying cell, 0 if no cell.
  - vtag (int array) contains the id of occupying vessel, 0 if no vessel.
- NOD: structure with data related to nodes. Arrays contain $M^2$ elements.
  - .fx, .fy: (float arrays) contain x and y component of the force that it is exerted by cells in this node.
  - .ux, .uy: (float arrays) contain x and y component of the displacement that it is suffered by this node.
  - .restrictx, restricty: (boolean arrays) contain 1 if there is a nodal restrictions in x or y direction, 0 otherwise.
- CEL: structure with data related to cells. Arrays contain a maximum of $10 \cdot (M-1)$ elements, but it is only filled with non-zero values up to the number of cells at that moment.
  - .siz: (int array) cell's size (in pixels).
  - .peri: (int array) cell's perimeter (in pixels).
  - .tip: (int array) position in the grid of the cell's pixel that is closer to the hypoxic area.

- .tail: (int array) position in the grid of the cell's pixel that is further to the hypoxic area.
- .vegf: (float array) concentration of VEGF associated to the cell (measured in bottom left grid point of the pixel selected in .tip).
- .pos: (int array) cell's type. 1 if it is a tip cell, 2 if it is a proliferating stalk cell and 3 if it is a non-proliferating stalk cell.
- .tshybrid: (boolean array) 1 if the cell has hybrid phenotype, 0 otherwise.
- .age: (int array) cell's age (in Monte Carlo Time Step, MCTS).
- .vess: (int array) id of the vessel that the cell belongs.

- NDJ: structure with the amount of cells proteins of signaling processes. Arrays contain a maximum of $10 \cdot (M-1)$ elements, but it is only filled with non-zero values up to the number of cells at that moment.
  - .N: (float array) amount of Notch in the cell.
  - .D: (float array) amount of Delta in the cell.
  - .J: (float array) amount of Jagged in the cell.
  - .I: (float array) amount of NICD in the cell.
  - .Vr: (float array) amount of VEGFR2 in the cell.
  - .V: (float array) amount of active VEGF in the cell.

- VES: structure with data related to vessels. Arrays contain a maximum of $M-1$ elements, but it is only filled with non-zero values up to the number of vessels at that moment.
  - .tiptag: (int array) id of the tip cell of the vessel.
  - .tipvox: (int array) id of the pixel stored on .tip (CEL) of the tip cell.
  - .proltag: (int array) id of proliferating stalk cell of the vessel.
  - .birth: (int array) MCTS when vessel was born.
  - .death: (int array) MCTS when vessel was died.
  - .isactive: (boolean array) 1 if the vessel is active, 0 otherwise.

- .branch: (int array) number of MCTS in which the new vessel has to maintain the direction or, after that, the id of the cell that could be a new tip cell of a new sprout.
- .bx, by: (float arrays) x and y coordinates of the branching direction during the number of MCTS that the vessel should follow it, 0.0 otherwise.
- .ncell: (int array) number of cells in the vessel.
- .parenttag: (int array) id of the vessel from which it branched, 0 if it is a initial sprout.
- .ndescen: (int array) number of sprouts that branched off the vessel.

- contact_perimeter: int array of size $10(M-1) \cdot 10(M-1)$. It is set as a matrix in which each row and column is the cell with the id of the row or column number. $(i, j)$ position stores the number of pixels shared by cell $i$ and cell $j$. $(i, i)$ position stores the number of perimeter pixels of cell $i$. $(i, 0)$ position stores the number of neighbors of cell $i$. $(0, i)$ position stores the number of perimeter pixels occupied by neighbors.
- V: int array of size $M \cdot M$. It stores the value of VEGF concentration on each node.

These structures and arrays have a copy in the CPU and another copy in the GPU. The main modules of the code work with the copy in the GPU and the copy in the CPU is updated to generate the output files.

3. Initialization of random numbers and structures

The CUDA Random Number Generation library (cuRAND) and the Mersenne twister algorithm are used in this code. After the declaration of all the variables needed, the Mersenne twister algorithm is initialized.

After that, all structures defined above are initialize at zero in CPU and copied to GPU (functions in `init.cpp` file and `init.cu`). At this point, we have two options:

- init_MCTS = 0, i.e., starts a new simulation. In function of the number of initial sprouts we have chosen, structures are modified to set new cells. Two kernels (kernel is function that runs on the GPU) are used to do that (kernels in `init.cu`). The first one assign one thread to one pixel and modify CEL and VOX structures, placing equispaced cells of one pixel sized. The second one assign one thread to one initial sprout and modify VES structures, filling data of the vessel.

- init_MCTS $\neq 0$, i.e., continues other simulation. In this case, data from output files is loaded in the CPU copy of the structures(functions in `read.cpp` file and `init.cu`).

4. Output files with used parameters and initial structures

In order to have registered the used parameters and the initial state of the simulation, output files with .out extension are generated.

`parameters.out` is a file with the name, value and description of the parameters set in `def.h`.

The data of structures is stored in the following files, where `X` is the value of init_MCTS:

- `ctagsX.out`: stores a $(M-1) \times (M-1)$ matrix with .ctag of VOX values.
- `vtagsX.out`: stores a $(M-1) \times (M-1)$ matrix with .vtag of VOX values.
- `dcnsX.out`: stores a matrix of size (cells number) $\times$ 15. Each row is a cell and each column is an item of the CEL (the first nine) and NDJ (the last six) structures, ordered as follows: .siz, .peri, .tip, .tail, .pos, .tshybrid, .age, .vess, .vegf, .N, .D, .J, .I, .V, Vr.
- `dvX.out`: stores a matrix of size (number of initial sprouts) $\times$ 12. Each row is a vessel and each column is an item of the VES structure, ordered as follows: .tiptag, .tipvox, .proltag, .birth, .death, .isactive, .branch, .bx, .by, .ncell, .parenttag, .ndescen.
- `periX.out`: stores a $10(M-1) \times 10(M-1)$ matrix with the contact_perimeter values.

5. Arrangement of ECM strains and displacements

In this module, the necessary preparations are made to calculate ECM strains and displacements, given by elasticity problem (9) of our manuscript. The matrix $K$ will remain fixed throughout the MCTS, therefore it is calculated once before starting. $K$ is a sparse matrix so the problem is needed to be solved with an appropriate method, like the Preconditioned Conjugate Gradient (PCG) method using ILU decomposition. The algorithm is provided by NVIDIA corporation and it is implemented on the GPU using CUBLAS and CUSPARSE libraries.

The following steps are performed in this module:

1. Set forces made by cells and forces in the boundary nodes. Set restrictions in the boundary nodes. These two actions are carried out in a single kernel that assign one thread to one node (kernel in `init.cu`).

2. Set element stiffness matrices $K_e$ (function in `FE_local.cpp`).

3. Assemble the global stiffness matrix $K$ from all element stiffness matrices (function in `FE_assemblyl.cp` [illegible]

4. Keep track of degree of freedom (DOF) of $K$ in an array. DOFs that are restricted get a -1, while the remaining DOFs get a number from 0 upwards (function in `FE_assemblyl.cpp`).

5. Reduce $K$ removing the row and column from K for all DOFs with a -1 (function in `FE_assemblyl.cpp`).

6. Prepare $K$ matrix in Compressed Sparse Row (CSR) format, necessary condition to use the PCG algorithm (function in `FE_assemblyl.cpp`). The steps 2 to 5 are functions of the supplementary material of Van Oers `et al.` [4]. However, we solved the system $Ku = f$ using parallel computing, so that $K$ matrix must be adapted. We have made this adaptation through a complex function that converts the output format of $K$ from Van Oers `et al.` [4] code to CSR format. $K$ is decomposed into three arrays:

   - $Kval$ is an array of size $10 \cdot 2 \cdot M \cdot M$. This array contains the non zero values of matrix $K$.

   - $Kcol$ is an array of size $10 \cdot 2 \cdot M \cdot M$. It contains the index of the column of corresponding nonzero value written in $Kval$.

   - $Krow$ is an array of size $N_{FE} + 1$, where $N_{FE}$ is the number of unrestricted DOFs. It contains the cumulative number of non-zero values in rows of $K$, starting with 0 in the first gap.

7. Prepare CUBLAS and CUSPARSE data:

   (a) Create CUBLAS context.

   (b) Create CUSPARSE context.

   (c) Description of the K matrix.

(d) Define the properties of the matrix.

(e) Create the analysis info object for the K matrix.

(f) Perform the analysis for the Non-Transpose case.

(g) Copy K data to ILU0 vals as input.

(h) Generate the Incomplete LU factor H for the matrix K using cudsparseScsrilu0.

(i) Create info objects for the ILU0 preconditioner.

6. Arrangement of VEGF concentration

In this module, the necessary preparations are made to calculate the concentration of VEGF, given by initial-boundary value problem (5) - (7) of our manuscript. Firstly, we introduce the following scalings in order to nondimensionalize:

$$\tilde{C} = \frac{C}{[C]},\ \tilde{t} = \frac{t}{[t]},\ \tilde{x} = \frac{x}{[x]},\ \tilde{y} = \frac{y}{[x]},\ \tilde{G} = \frac{G}{[G]}$$

where $[C]$ , $[t]$ , $[x]$ and $[G]$ are non-zero parameters. Substituting these new variables into the PDE, we obtain:

$$\frac{[C]}{[t]}\frac{\partial \tilde{C}}{\partial \tilde{t}} = \frac{D_f\,[C]}{[x]^2}\left(\frac{\partial^2 \tilde{C}}{\partial \tilde{x}^2} + \frac{\partial^2 \tilde{C}}{\partial \tilde{y}^2}\right) - \nu\,[C]\,\tilde{C} - [G]\,\tilde{G} \;\Leftrightarrow\; \left(\text{dividing by } \frac{D_f[C]}{[x]^2}\right)$$

$$\frac{[x]^2}{D_f\,[t]}\frac{\partial \tilde{C}}{\partial \tilde{t}} = \left(\frac{\partial^2 \tilde{C}}{\partial \tilde{x}^2} + \frac{\partial^2 \tilde{C}}{\partial \tilde{y}^2}\right) - \frac{[x]^2\,\nu}{D_f}\tilde{C} - \frac{[G]\,[x]^2}{D_f\,[C]}\tilde{G}$$

Regarding to the cell binding term, choosing $[G] = \frac{D_f[C]}{[x]^2}$ leads to have two terms which are $\mathcal{O}(1)$, cell binding term and diffusion term, and the PDE obtained is:

$$\frac{[x]^2}{D_f\,[t]}\frac{\partial \tilde{C}}{\partial \tilde{t}} = \triangle \tilde{C} - \frac{[x]^2\,\nu}{D_f}\tilde{C} - \tilde{G}$$

The value of $\frac{[x]^2}{D_f[t]}$ and of $\frac{[x]^2\nu}{D_f}$ depend on the characteristic time scale, equal to 1 MCS, and the characteristic length scale, on the order of the length of a side of 1 voxel. The value of $[C]$ depends on $S$ and the value of $[G]$ could be known with these data. Regarding to the equivalence between 1 MCS and the real time units, we use the time of the experiments in the work of Sugihara *et al.* [5] to calibrate Monte Carlo steps. In this work, the branch elongation takes 36 hours to grow 135 $\mu$m, so it would take 132 hours to grow 495 $\mu$m. If our simulations take 3001 MCS approximately, 1 MCS is 0.044 hours.

$$[t] = 1 \text{ MCS} \simeq 0.044 \text{ h}, \; [x]^2 = \left(\frac{L}{M-1}\right)^2 = \left(\frac{0.495 \text{ mm}}{600}\right)^2 = 0.6806 \; \mu\text{m}^2 = h^2$$

$$[C] = \frac{1}{3}S = 0.005 \text{ pg}/\mu\text{m}^2 \text{ due to the triangle centroid, so}$$

$$[G] = \frac{D_f\,[C]}{[x]^2} = \frac{3.6 \times 10^4 \; \mu\text{m}^2/\text{h} \times 0.005 \text{ pg}/\mu\text{m}^2}{0.6806 \; \mu\text{m}^2} = 264.47 \text{ pg}/(\mu\text{m}^2 \text{ h})$$

Therefore,

$$\frac{[x]^2}{D_f\,[t]} = \frac{0.6806 \; \mu\text{m}^2}{3.6 \times 10^4 \; \mu\text{m}^2/\text{h} \times 1\text{h}} = 4.2967 \times 10^{-4} \ll 1$$

$$\frac{[x]^2\,\nu}{D_f} = \frac{0.6806 \; \mu\text{m}^2 \times 0.6498 \text{ h}^{-1}}{3.6 \times 10^4 \; \mu\text{m}^2/\text{h}} = 1.2285 \times 10^{-5} \ll 1$$

The resulting factors are much less than one, so the solution of

$$0 = \triangle\tilde{C}(\tilde{x}, \tilde{y}, \tilde{t}) - \tilde{G}(\tilde{x}, \tilde{y}, \tilde{C}), \; (\tilde{x}, \tilde{y}) \in \Omega, \; \tilde{t} > 0$$

$$C(0, \tilde{y}, \tilde{t}) = 0, \; C(M-1, \tilde{y}, \tilde{t}) = \tilde{S} = \frac{S}{[C]}, \; C(\tilde{x}, 0, \tilde{t}) = \frac{\tilde{S}}{M-1}\tilde{x} = C(\tilde{x}, M-1, \tilde{t}), \; (\tilde{x}, \tilde{y}) \in \Omega, \; \tilde{t} > 0$$
$$C(\tilde{x}, \tilde{y}, 0) = 0, \; (\tilde{x}, \tilde{y}) \in \Omega$$

can be used to approximate the VEGF field in our model. This equation can be solved with the Finite Difference Method (FDM) using five-point stencil. [2].

But, first of all and in order to simplify the notation, dimensionless parameters and functions will be used without tildes. The notation $\nabla^2$ is more suitable for the laplacian operator; the symbol $\triangle$ would lead to confusion in numerical work where $\triangle x$ and $\triangle y$ are used for grid spacing. To sum up, we obtain the following Poisson problem:

$$\begin{aligned}
\text{PDE:} \quad & \nabla^2 C(x, y, t) = G(x, y, C), \; 0 < x < M-1, \; 0 < y < M-1, \; t > 0 \\
\text{BC:} \quad & C(0, y, t) = 0, \; C(M-1, y, t) = S, \; 0 < y < M-1, \; t > 0 \\
& C(x, 0, t) = \tfrac{S}{M-1}x = C(x, M-1, t), \; 0 < x < M-1, \; t > 0 \\
\text{IC:} \quad & C(x, y, 0) = 0, \; 0 < x < M-1, \; 0 < y < M-1.
\end{aligned} \tag{0.1}$$

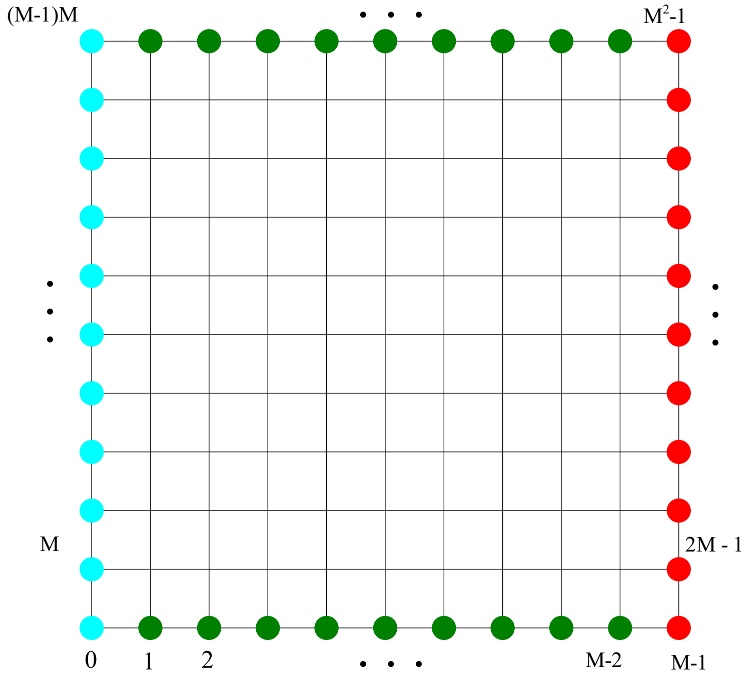


Figure 2: Grid scheme.

Let $C_{ij}$ represent an approximation to $C(x_i, y_j, t)$, where $(x_i, y_j)$ have been described in domain section. Note that time variable is not been taken into account because the solution of the PDE of (0.1) on each MCS is not time-dependent. To discretize the PDE of (0.1), centered finite differences have been used for $x-$ and $y-$derivatives, which gives:

$$\begin{gathered}\tfrac{1}{h^2}\left(C_{i-1,j} - 2C_{ij} + C_{i+1,j}\right) + \tfrac{1}{h^2}\left(C_{i,j-1} - 2C_{ij} + C_{i,j+1}\right) = G_{ij} \Leftrightarrow \\ \left(C_{i-1,j} + C_{i,j-1} - 4C_{ij} + C_{i+1,j} + C_{i,j+1}\right) = h^2 G_{ij}\end{gathered} \tag{0.2}$$

where $G_{ij} = G(x_i, y_j)$ (we know exactly $G$ function). The equation (0.2) changes in function of the point it is focus on and some of them have boundary conditions, so the following equations are written distinguishing each case:

- If $j = 0$ or $j = M - 1$ and $\forall i$ (green nodes in figure 2 (b)),

$$C_{i,0} = C_{i,M-1} = \frac{S}{M-1} i$$

- If $i = 0$ and $\forall j$ (blue nodes in figure 2 (b)),

$$C_{0j} = 0,$$

- If $i = M - 1$ and $\forall j$ (red nodes in figure 2 (b)),

$$C_{M-1j} = S,$$

- If $i = 1$ and:
  - $j = 1$

$$(-4C_{1,1} + C_{1,2} + C_{2,1}) = h^2 G_{1,1} - \frac{S}{M-1}, \tag{0.3}$$

  - $j = M - 2$,

$$(C_{1,M-3} - 4C_{1,M-2} + C_{2,M-2}) = h^2 G_{1,M-2} - \frac{S}{M-1}, \tag{0.4}$$

  - $2 \leq j \leq M - 3$,

$$(C_{1,j-1} - 4C_{1,j} + C_{1,j+1} + C_{2,j}) = h^2 G_{1,j}, \tag{0.5}$$

- If $i = M - 2$ and:

- $j = 1$

$$(C_{M-3,1} - 4C_{M-2,1} + C_{M-2,2}) = h^2 G_{M-2,1} - \frac{S}{M-1}(M-2) - S, \tag{0.6}$$

- $j = M-2$,

$$(C_{M-3,M-2} + C_{M-2,M-3} - 4C_{M-2,M-2}) = h^2 G_{M-2,M-2} - \frac{S}{M-1}(M-2) - S, \tag{0.7}$$

- $2 \leq j \leq M-3$,

$$(C_{M-3,j} + C_{M-2,j-1} - 4C_{M-2,j} + C_{M-2,j+1}) = h^2 G_{M-2,j} - S, \tag{0.8}$$

- If $j = 1$ and $2 \leq i \leq M-3$

$$(C_{i-1,1} - 4C_{i,1} + C_{i,2} + C_{i+1,1}) = h^2 G_{i,1} - \frac{S}{M-1} i, \tag{0.9}$$

- If $j = M-2$ and $2 \leq i \leq M-3$

$$(C_{i-1,M-2} + C_{i,M-3} - 4C_{i,M-2} + C_{i+1,M-2}) = h^2 G_{i,M-2} - \frac{S}{M-1} i, \tag{0.10}$$

- If $2 \leq i \leq M-3$ and $2 \leq j \leq M-3$,

$$(C_{i-1,j} + C_{i,j-1} - 4C_{ij} + C_{i,j+1} + C_{i+1,j}) = h^2 G_{ij}, \tag{0.11}$$

The above equations can be collected together into a matrix equation $Ax = R$. The coloured nodes of the figure 2 (b) are not considered since their values are already known. So, the equations to take into account are from (0.3) to (0.11). $A$ is a matrix of size $N_{FD}^2$, with $N_{FD} = (M-3)^2$ that contains the factors that appear multiplying $C_{ij}$ terms, i.e. 0, 1 and $-4$. $x$ is an array of size $N_{FD}$ that contains the values for $C_{ij}$ terms that satisfy the matrix form equation. The first time we calculate $x$, it is initialized at zero, but in future Monte Carlo time steps we use the solution calculated in the previous MCTS. $R$ is an array of size $N_{FD}$ and it is formed by the right - hand side (RHS) of the previous equations.

$A$ is a sparse matrix so the system is needed to be solved with an appropriate method, like the Preconditioned Conjugate Gradient (PCG) method using ILU decomposition. The algorithm is provided by CUDA and uses a Compressed Sparse Row (CSR) format, so we have decomposed $A$ into three arrays:

- *Aval* is an array of size $nz_{FD} = 5N_{FD} - 4\sqrt{N_{FD}}$. $nz_{FD}$ is the number of non zero values of $A$, i.e. five - point finite difference scheme minus number of frontier nodes per boundary times number of boundaries. This array contains the non zero values of matrix $A$.
- *Acol* is an array of size $nz_{FD}$. It contains the index of the column of corresponding nonzero value writen in *Aval*.
- *Arow* is an array of size $N_{FD} + 1$. It contains the cumulative number of non-zero values in rows of $A$, starting with 0 in the first gap.

These three arrays, plus $x$ and $R$, are needed to use the mentioned algorithm. After that, the $x$ array is included in a bigger one $V$, which also contains the values of VEGF concentration at the boundaries.

The following steps are performed in this module:

1. Initialize $x$ and $R$.
2. Define $A$ matrix in CSR format.
3. Prepare CUBLAS and CUSPARSE data:
    (a) Create CUBLAS context.
    (b) Create CUSPARSE context.
    (c) Description of the A matrix.
    (d) Define the properties of the matrix.
    (e) Create the analysis info object for the A matrix.
    (f) Perform the analysis for the Non-Transpose case.
    (g) Copy A data to ILU0 vals as input.
    (h) Generate the Incomplete LU factor H for the matrix A using cudsparseScsrilu0.
    (i) Create info objects for the ILU0 preconditioner.
4. Solve $Ax = R$ system using PCG method.
5. Include array $x$ in $V$. This takes place in a kernel where is assigned one thread to one node (kernel in `chemotaxis.cu`).

6. Associate to each cell the value of VEGF concentration on the bottom left grid point of the pixel selected in .tip of CEL structure, i.e., update .vegf of CEL structure. This takes place in a kernel where is assigned one thread to one cell (kernel in `chemotaxis.cu`).

7 & 15. Output files with structures

These output files are used to save data about the simulation every certain MCTS and then to be able to visualize some data or make statistics. Files are generated with an .out extension; they store data of structures. The frequency at which these data are stored can be changed according to interest of the simulation, but normally it is every 50 MCTS. But before generating the files, the copy in the CPU is updated with the new data of the structures that are in the GPU.

The generated files in MCTS `X` are:

- `ctagsX.out`: stores a $(M-1)\times(M-1)$ matrix with .ctag of VOX values.
- `vtagsX.out`: stores a $(M-1)\times(M-1)$ matrix with .vtag of VOX values.
- `dcnsX.out`: stores a matrix of size (cells number) $\times$ 15. Each row is a cell and each column is an item of the CEL (the first nine) and NDJ (the last six) structures, ordered as follows: .siz, .peri, .tip, .tail, .pos, .tshybrid, .age, .vess, .vegf, .N, .D, .J, .I, .V, Vr.
- `dvX.out`: stores a matrix of size (number of initial sprouts) $\times$ 12. Each row is a vessel and each column is an item of the VES structure, ordered as follows: .tiptag, .tipvox, .proltag, .birth, .death, .isactive, .branch, .bx, .by, .ncell, .parenttag, .ndescen.
- `pstrain.out`: stores a matrix of size $(M\cdot M)\times$ 6. Each row is a node and columns are: .fx, .fy, larger eigenvalue, first component of the eigenvector of this eigenvalue, second component of the eigenvector of this eigenvalue, the other eigenvalue.
- `periX.out`: stores a $10(M-1)\times 10(M-1)$ matrix with the contact_perimeter values.
- `VEGF.out`: stores a $M\times M$ matrix with values of V.

8. Branching module

The following steps are performed in this module (kernels in `branching.cu`):

1. Kernel that calculates the direction of branching of each possible new tip cells. This kernel assigns one thread to one pixel.

2. Kernel that calculates the probability of branching taking into account the direction and their neighbors. This kernel assigns one thread to one vessel.

3. Kernel that updates VOX structure with new tip cells, in the case where branching has been accepted. This kernel assigns one thread to one pixel.

4. Kernel that updates CEL and VES structure. This kernel assigns one thread to one vessel.

5. Kernel that checks if one of these new tip cells needs to jump in the direction of branching and update CEL and VES structures, if necessary. This kernel assigns one thread to one vessel.

6. Kernel that updates VOX structure after cell's jumping, if necessary. This kernel assigns one thread to one pixel.

For more details about the branching process, look up the main text.

9. Cell proliferation module

The following steps are performed in this module (kernels in `proliferation.cu`):

1. Kernel that builds an array with the labels of the cell that can proliferate for each vessel. This kernel assigns one thread to one vessel.

2. Kernel that checks if cells in the built array meet certain requirements to proliferate. *nprol* is the number of cells that have met the requirements. This kernel assigns one thread to one vessel.

3. Kernel that prepares data arrays for K-means algorithm. This kernel runs *nprol* times.

4. K-means algorithm is used to form two groups of pixels of each cell. After that, every structure and contact_perimeter array are updated with the new data of these two cell that have formed through the groups of pixels. The two kernels involved run sequentially in a loop, *nprol* times, one for each cell.

For more details about the cells proliferation process, consult our manuscript.

10. VEGF concentration module

The following steps are performed in this module (kernels in `chemotaxis.cu`):

1. Kernel that calculates right-hand side $R$ of the system $Ax = R$ (see 6. Arrangement of VEGF concentration). This kernel assigns one thread to one node.
2. Solve $Ax = R$ system using PCG method where $x$ is the solution calculated in the last MCTS.
3. Kernel that includes the array $x$ in $V$. This kernel assigns one thread to one node.
4. Kernel that associates to each cell the value of VEGF concentration on the bottom left grid point of the pixel selected in .tip of CEL structure, i.e., update .vegf of CEL structure. This kernel assigns one thread to one cell.

11. Signaling processes module

In this module, the nondimensionalized version of the system of Ordinary Differential Equations (ODEs), Eqs. (14) - (19) of the main text, is numerically solved.

This system is solved with the explicit Euler method each 10 MCS, due to the difficulty of parallelize this module. According to Boareto *et al.* [1], the role of tip and stalk cells could change every two hours, so that updating parameters values of the proteins involved each 0.44 hours is enough.

On each step of time, a system of six ODEs musts to be solved on each cell. This is done in a kernel that assigns one thread to one cell and calculates approximate solution with the explicit Euler method to the cell. Then, the approximate solution is incorporated to NS structure through a kernel that assign also one thread to one cell. The maximum number of steps has been chosen takes into account the convergence of the Euler method. Other numerical methods have been taken into account but have been discarded because they required more computational time and the error was approximately the same for the time step size we use.

12. ECM strains and displacements module

The following steps are performed in this module:

1. Kernel that restarts forces in NOD structure. This kernel assigns one thread to one node (kernel in `cellforces.cu`).

2. Kernel that calculates forces on each node and writes them in NOD structure. This kernel assigns one thread to one node (kernel in `cellforces.cu`).

3. Kernel that copies displacements calculated on the last MCTS from NOD structure in an array $u$. This kernel assigns one thread to one node (kernel in `FE_nodes2dofs.cu`).

4. Kernel that copies calculated forces in an array $f$. This kernel assigns one thread to one node (kernel in `FE_nodes2dofs.cu`).

5. Solve $Ku = f$ system using PCG method.

6. Kernel that copies the recalculated $u$ in NOD structure. This kernel assigns one thread to one node (kernel in `FE_nodes2dofs.cu`).

13. CPM and anastomosis module

The following steps are performed in this module (kernels in `cellmoves.cu`):

1. Generate three arrays of size $(M-1)\cdot(M-1)$ with:

    - Id of randomly selected pixels.
    - Id of random neighbors of randomly selected pixels.
    - Random float numbers between 0 and 1 used to calculate the Boltzmann probability factor in the Metropolis algorithm.

2. We divided the grid in boxes of pixels in order to compute each one in parallel. Therefore we need to know to which box each randomly selected pixel belongs. It is done by a kernel that assigns one thread to one pixel.

3. The kernel in charge of the CPM and the anastomosis assigns one thread to one box of pixels. In parallel on each box, a loop runs through each randomly selected pixel that it contains. The steps that are followed in this kernel are:

(a) In the case of the selected pixel belongs to a expanding cell of a new sprout, check if the direction of expansion is similar to the branching direction.

(b) Check if a cell will break in two, if necessary.

(c) Calculate the variation of energy if the copy is made with the Hamiltonian $H$ described in our manuscript.

(d) Calculate the Boltzmann probability and check, using a random float number previously calculated, whether, according to the Metropolis rule, the copy is approved.

(e) If the copy is approved, update structures and contact_perimeter array.

(f) Check if anastomosis has occurred between two vessels and one of them becomes inactive.

14. Update features module

The following steps are performed in this module (kernels in `proliferation.cu`):

1. Kernel that updates the age of cells, i.e., .age of CEL structure. This kernel assigns one thread to one cell.

2. Kernel that updates the size of cells and pixels that are closer and further to the hypoxic area of cells, i.e., .siz, .tip and .tail of CEL structure. This kernel assigns one thread to one pixel.

3. Kernel that updates the number of MCTS that a sprout has to keep its branching direction, i.e., the incubation time of our manuscript. This kernel assigns one thread to one vessel.

4. Kernel that updates types of cells and selects possible new tip cells of new sprouts. this kernel assigns one thread to one pixel.

⋆ Compiling and IDE

We have used the integrated development environment (IDE) Microsoft Visual Studio to edit the code, to compile multiple source files (including .cu files and CUDA libraries) and build the executable file.

⋆ Hardware and computation time

The computation time of each simulation in a computer with Intel(R) Core(TM) i7-7700K CPU @4.20 GHz processor, 64.0 GB RAM and NVIDIA GeForce GTX 1080 graphics card is about 4 hours.
